# Idiosyncratic epistasis leads to global fitness-correlated trends

**DOI:** 10.1101/2021.09.22.461382

**Authors:** Christopher W. Bakerlee, Alex N. Nguyen Ba, Yekaterina Shulgina, Jose I. Rojas Echenique, Michael M. Desai

## Abstract

Epistasis can dramatically affect evolutionary trajectories. In recent decades, protein-level fitness landscapes have revealed extensive idiosyncratic epistasis among specific mutations. In contrast, other work has found ubiquitous and apparently non-specific patterns of global diminishing-returns and increasing-costs epistasis among mutations across the genome. Here, we use a hierarchical CRISPR gene drive system to construct all combinations of 10 missense mutations from across the genome in budding yeast, and measure their fitness in six environments. We show that the resulting fitness landscapes exhibit global fitness-correlated trends, but that these trends emerge from specific idiosyncratic interactions. This provides the first experimental validation of recent theoretical work that has argued that fitness-correlated trends can emerge as the generic consequence of idiosyncratic epistasis.

**One-Sentence Summary:** A genome-spanning fitness landscape reveals how idiosyncratic genetic interactions lead to global epistatic patterns.

## Main Text

Epistatic interactions have important consequences for the design and evolution of genetic systems (*1–3*). Significant work in recent decades has studied these interactions by measuring empirical fitness landscapes, most often at “shallow” depth for genome-scale studies (e.g., by quantifying pairwise but not higher order epistasis between all gene deletions or mutations) or at “narrow” breadth (such as complete landscapes at the scale of small select regions in single genes, for example by quantifying all orders of epistatic interactions among few amino acid residues) (*4–18*). These studies have often found many epistatic interactions among specific mutations at both lower (i.e., among few mutations) and higher orders (i.e., among many mutations). These reflect particular biological and physical interactions among the mutations involved; following recent work (*19, 20*) we refer to them as “idiosyncratic” epistasis, as they involve the specific details of these mutations. Overall, this body of work has highlighted the potential for epistasis to create historical contingency that tightly constrains the distribution of adaptive trajectories accessible to natural selection.

In contrast, other work examining adaptive trajectories that implicate loci across the genome has found patterns of apparently “global” epistasis, in which the fitness effect of a mutation varies systematically with the fitness of the genetic background on which it occurs (*21–28*). Typically, this manifests as either diminishing returns for beneficial mutations or increasing costs for deleterious mutations, with mutations having a less positive or more negative effect on fitter backgrounds. These consistent patterns of global epistasis may give rise to the dominant evolutionary trend of declining adaptability, and in contrast to the complexity of idiosyncratic interactions, they suggest that historical contingency could be less critical in constraining adaptive trajectories (*29*).

Despite their importance, these dual descriptions of epistasis have not been satisfactorily unified. In one view, global epistasis results from non-specific fitness-mediated interactions among mutations (*24*). Such interactions may for example emerge from the topology of metabolic networks, which generates overall patterns of diminishing returns and increasing costs that eclipse the specific details of idiosyncratic interactions (*30*). In contrast, other recent theoretical work has proposed an alternative view, hypothesizing that apparent fitness-mediated epistasis can instead emerge as the generic consequence of idiosyncratic interactions, provided they are sufficiently numerous and widespread (*19*, *20*). These two models have substantially different implications for the structure of fitness landscapes, which in turn influence our expectations of the repeatability and predictability of evolution and of the effect of chance and contingency on adaptation at both the genotypic and phenotypic level. Thus, this dichotomy plays a central role in understanding of how epistasis affects evolutionary dynamics.

Thus far, however, empirical work has been unable to distinguish between these perspectives. The key difficulty is that testing these ideas requires both depth and breadth: we must analyze landscapes involving enough loci that we sample idiosyncratic interactions that can potentially give rise to overall fitness-mediated trends, and we must survey possible combinations of these mutations at sufficient depth to quantify the role of higher-order interactions (including potential “global” non-specific fitness-mediated interactions). Importantly, larger landscapes are also necessary to reduce the influence of measurement error on the inference of epistasis and analysis of fitness-correlated trends (see SI section 6.3). Achieving this depth and breadth is technically challenging, because it requires us to synchronize many mutations across the genome.

Here, we overcome this challenge by developing a novel method that exploits Cre-Lox recombination to create a combinatorially expanding CRISPR guide-RNA (gRNA) array in *Saccharomyces cerevisiae,* which allows us to iteratively generate mutations at distant loci via a gene drive mechanism (Fig. 1A). Briefly, strains of opposite mating type containing inducible Cre recombinase and *SpCas9* genes are mutated at one of two loci (*A* or *B*), and DNA encoding guide-RNAs (gRNAs) specific to the wild-type alleles at these loci are integrated into their genomes (Fig. S1). After mating to produce a diploid heterozygous at *A* and *B,* we induce a gene drive to make the loci homozygous. This begins with expressing Cas9 and generating gRNA-directed double-strand breaks at the wild-type *A* and *B* alleles. These breaks are then repaired by the mutated regions of homologous chromosomes, making the diploid homozygous at these loci with at least 95% efficiency. Simultaneously, we express Cre to induce recombination that brings gRNAs into physical proximity on the same chromosome by way of flanking Lox sites, in a strategy similar to that described previously (*31*) (Fig. 1B). We then sporulate diploids and select haploids bearing the linked gRNAs from both parents. In parallel, we carry out this process with “pseudo-WT” versions of these loci, which contain synonymous changes that abolish gRNA recognition, but lack the non-synonymous change of interest. This creates a set of four strains, with all possible genotypes at loci *A* and *B*. Concurrently, we create separate sets of four strains with all possible genotypes at other pairs of loci (e.g., *C* and *D*).

**Fig. 1.**
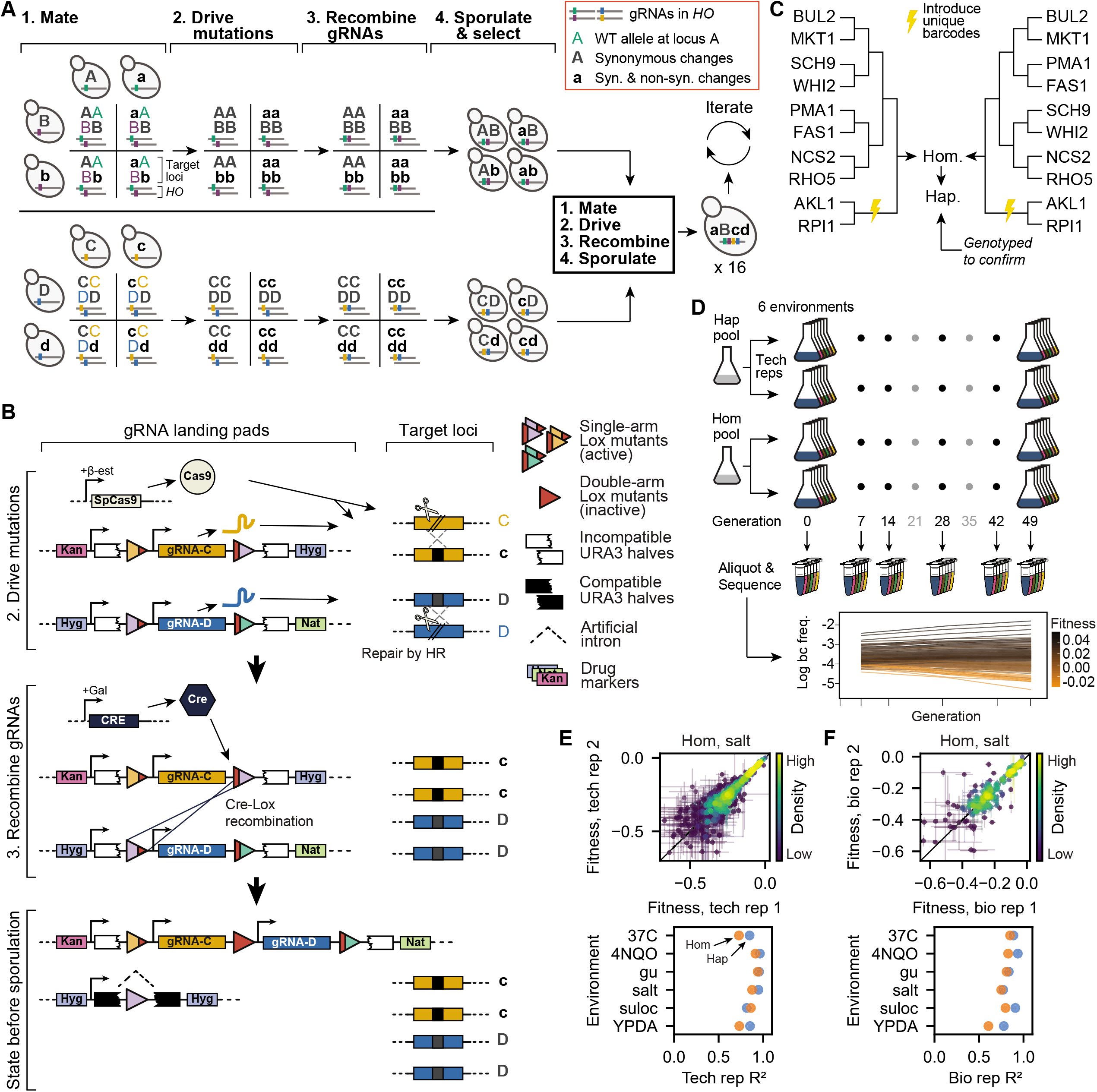
Recombining CRISPR-gene drive system. **(A)** Experimental design. Strains of opposite mating type carrying known mutations and corresponding guide-RNAs (gRNAs) mate to form heterozygous diploids. Cas9 expression “drives” these mutations, and site-specific recombination links gRNAs. Homozygous diploids are sporulated, haploids with linked gRNAs are selected, and the process repeats, incorporating exponentially increasing numbers of mutations. **(B)** Recombining gene drive system. gRNAs targeting heterozygotic loci are flanked by selection markers and two of three orthogonal Lox sites (colored triangles), which are inactivated through recombination (red triangles). Cas9 “drives” targeted mutations, while Cre-Lox recombination brings like markers to the same chromosome and activates a *URA3* gene interrupted by an artificial intron. Following sporulation, the chromosome with gRNAs is selected using the markers of interest while the other is counterselected using 5-FOA. **(C)** Cross design. A complete fitness landscape is produced in parallel by distinct cross designs that yield final homozygous diploids and haploids in biological replicates with unique DNA barcodes. **(D)** Bulk-fitness assays. Pooled strains are assayed in replicate for competitive fitness in several environments by sequencing barcodes to obtain strain frequencies over time. **(E)** Repeatability of technical replicate competitive fitness measurements. **(F)** Repeatability of biological replicate competitive fitness measurements.

By iterating this process, we can rapidly assemble an exponentially expanding, combinatorially complete genotype library. We mate separate sets of four genotypes bearing all combinations of mutations at two loci each in an all-against-all cross, drive their mutations, recombine their gRNAs, and sporulate to produce a 16-strain library bearing all 4-locus mutation combinations. Repeating these steps in a third cycle with two 4-locus libraries of opposite mating type yields a 256-strain 8-locus library, and a complete landscape of up to 16 mutations (2^16^ strains) can be constructed in just four cycles.

We sought to use this method to construct a complete fitness landscape that would shed light on the structure of epistasis: are fitness-correlated trends primarily the product of a global coupling of mutations via fitness, or do they emerge as the consequence of idiosyncratic epistasis? To do so, we surveyed studies of natural variation (e.g., (*32–36*)) and experimental evolution (e.g., (*37–39*)) to identify mutations relevant to adaptation in the laboratory strain. We selected a set of mutations that sample a wide range of cellular functions, such as membrane stress response, mitochondrial stability, and nutrient sensing. Our goal in making this choice was to maximize fitness variance while minimizing pathway-specific idiosyncratic interactions. We note that alternative choices of mutations, particularly if they were focused on a specific protein or pathway (or limited to those that accumulated along the line of descent in a single lineage), might exhibit very different patterns of epistasis, which would be characteristic of the particular details of that specific protein or pathway (or that specific adaptive trajectory). However, our goal here is to analyze potentially global patterns of epistasis among mutations across the genome that are relevant to adaptation and hence represent an overall fitness landscape for the laboratory strain.

We thus implemented our gene-drive system to construct a near-complete landscape spanning 10 missense mutations in 10 genes (including essential genes) on 8 chromosomes: *AKL1* (S176P), *BUL2* (L883F), *FAS1* (G588A), *MKT1* (D30G), *NCS2* (H71L), *PMA1* (S234C), *RHO5* (G10S), *RPI1* (E102D), *SCH9* (P220S), and *WHI2* (L262S) (Fig. 1C, Table S1). We found that a landscape of about this size is required to distinguish the two models (see SI section 6.3). Immediately before the final mating cycle, all strains were transformed with a unique DNA barcode next to the *LYS2* locus to enable high-throughput, sequencing-based competitive fitness assays (Fig. S2, S3). All strains in each replicate haploid library were genotyped at all 10 loci to confirm presence of the desired alleles (this also ensures presence in the diploid libraries). After excluding strains due to gene drive failure, 875 out of 1024 (85.4%) genotypes remained in at least one library (and 407 in both biological replicates). We also performed whole genome sequencing of 96 randomly selected strains to rule out pervasive aneuploidies or influential but spurious background mutations. One aneuploidy was identified, and 3 spurious background mutations were observed at >5% frequency. Subsequent analysis showed that these were unlikely to systematically influence our findings (Table S2, and SI section 5.1).

To obtain fitness landscapes, we conducted replicate bulk barcode-based fitness assays on both pooled haploid and homozygous diploid versions of the genotype library in 6 distinctly stressful media environments: YPD + 0.4% acetic acid, YPD + 6 mM guanidium chloride, YPD + 35 μM suloctidil, YPD @ 37°C, YPD + 0.8 M NaCl, and SD + 10 ng/mL 4NQO (Fig. 1D). For each of 7 days, pools were allowed 7 generations of growth, and aliquots were sampled and sequenced at the barcode locus at generations 7, 14, 28, 42, and 49. We estimated the relative fitness of each genotype from changes in barcode frequencies through time, achieving consistent measurements across technical and biological replicates (Fig. 1E,F, S4). From these data, we inferred the background-averaged additive and epistatic effects of each mutation and combination of mutations, respectively (using LASSO regularization, see SI).

We found that our six environments yield substantially different landscapes, as demonstrated by the relatively low between-environment correlations of genotype fitnesses (Fig. 2A), the additive effects of each mutation (Fig. 2B), and the pairwise interactions between them (Fig. 2C). While haploid and homozygous diploid landscapes were largely correlated, there were several notable exceptions, particularly in the suloctidil environment (Fig. 2A,B). And while some pairwise interactions remain roughly constant in strength, even as the corresponding additive effects vary considerably (e.g., *RHO5* and *WHI2*), most wax and wane across environments (Fig. 2C). Nevertheless, the overall contribution from different epistatic orders shows some similarities across ploidies and environments (the magnitudes do differ; Fig. 2D), with additive and pairwise terms explaining most of the variance in the data, third-order terms contributing minorly, and the remaining orders making little difference, consistent with earlier studies (*40*). Across all epistatic orders, inferred effects were highly skewed, with a small number of terms explaining disproportionate variance (Fig. 2E).

**Fig. 2.**
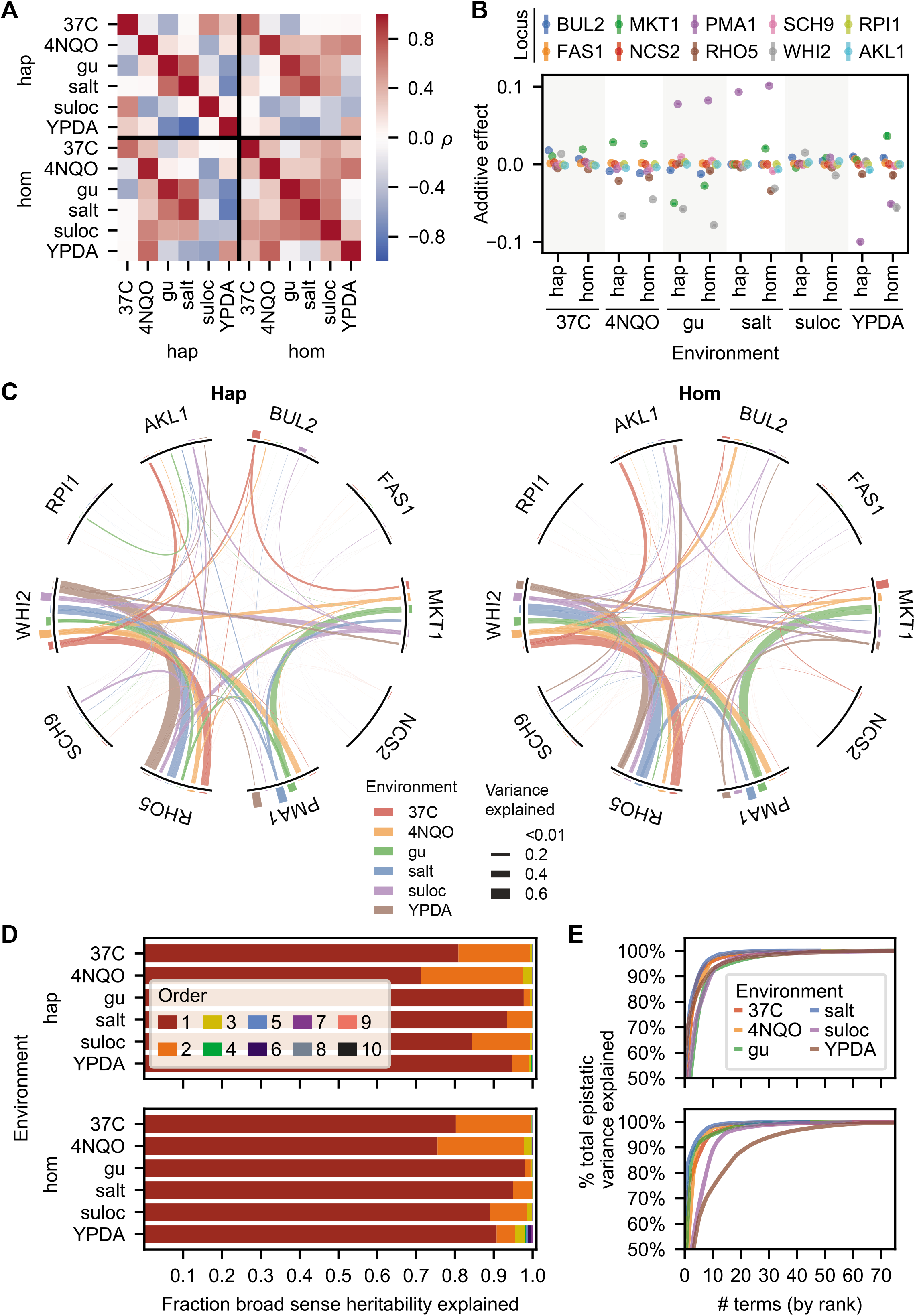
Fitness landscapes. **(A)** Correlation in observed fitness (upper right) and predicted fitness (from inferred model, lower left, see SI section 5.1) across ploidies and environments. **(B)** Background-averaged additive effect of each locus across ploidies and environments. Error bars represent 95% confidence intervals. **(C)** Background-averaged pairwise epistatic effects between loci across ploidies and environments. Weights of edges connecting loci represent the proportion of pairwise variance explained by each interaction. Heights of bars on the perimeter correspond to the proportion of additive variance explained by each locus in each environment. **(D)** Variance partitioning of broad-sense heritability from additive and epistatic orders across ploidies and environments. **(E)** Cumulative distribution of the epistatic variance explained by rank-ordered epistatic terms of all orders.

We next sought to investigate potential patterns of global fitness-mediated epistasis. To do so, for each locus in each ploidy and environment, we plotted the fitness of a genotype with the mutated allele, φ_Mut_, against the fitness of the same genotype with the WT allele, φ_WT_. A regression slope, *b*, different from 1 in these plots signifies a fitness-correlated trend (FCT) (Fig. 3A, left; see SI). We note that some previous work has instead plotted the fitness effect of a mutation, Δφ, as a function of background fitness φ_WT_. The advantage of our formulation here is that it does not privilege a specific allele as the “wild-type.” Instead, regression in our plots translates intuitively when reversing direction to treat the reversion as the mutation: *b*_rev_ = 1/*b*_orig_ by weighted-total least squares; see expanded discussion in the SI, Fig. S5-S8.

**Fig. 3.**
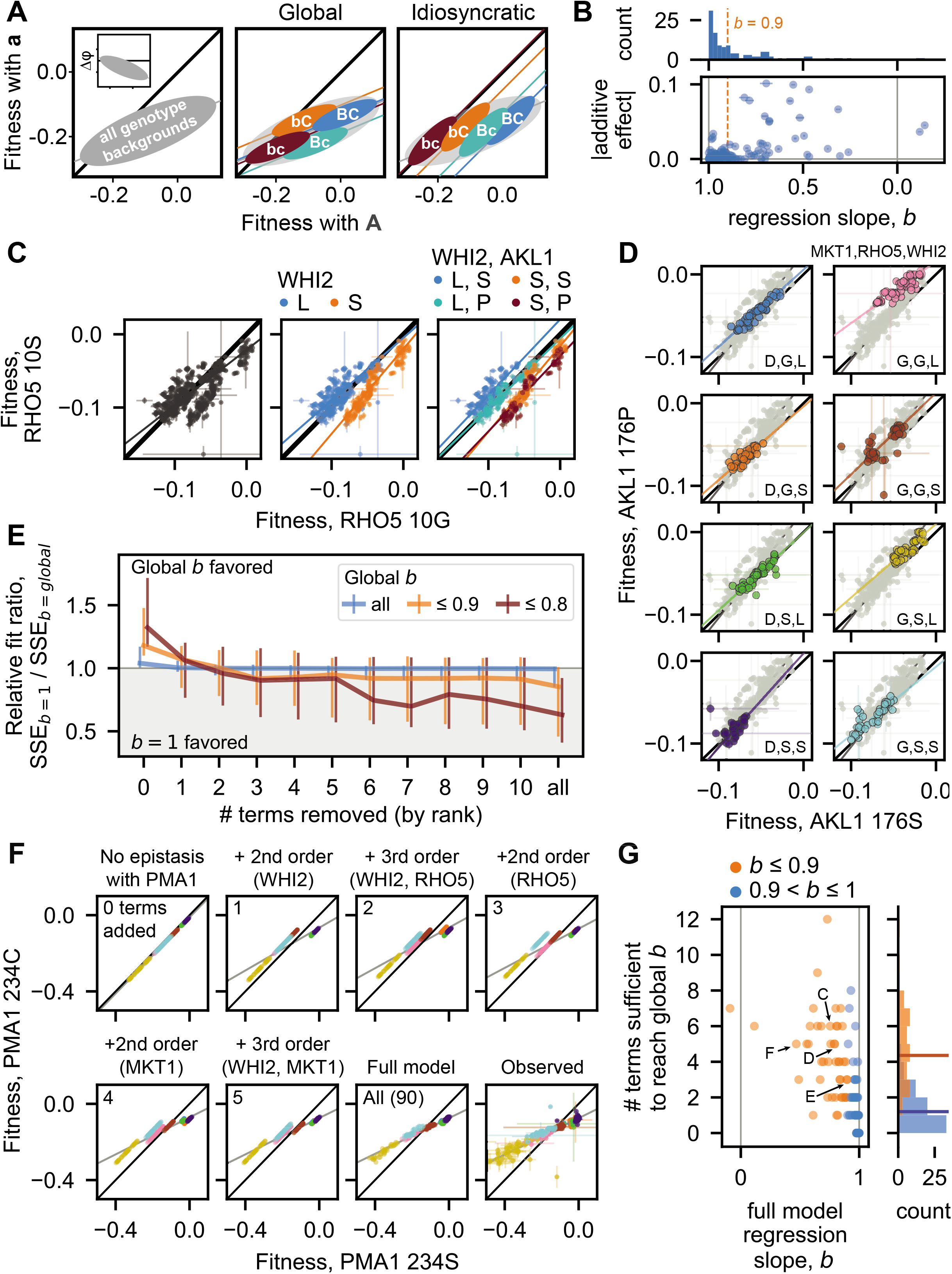
Fitness-correlated trends (FCTs). **(A)** Schematic contrasting how global or idiosyncratic epistasis could produce FCTs. Inset shows FCT analyzed as the effect of a mutation (Δφ) on backgrounds of different fitnesses. **(B)** Histogram and scatterplot of regression slopes, *b,* between φ_Mut_ and φ_WT_, and corresponding absolute additive effects of mutations. Polarity adopted such that *b* ≤ 1. Total error bar length is twice the standard error of the slope. **(C)** Fitness effect of *RHO5* mutation (G10S) (φMut versus φ_WT_) in all haploid backgrounds at 37°C (left) and partitioned by genotypes at *WHI2* (L262S) (middle) and *WHI2* and *AKL1* (S176P) (right). Initial SSE_*b*=1_ / SSE_*b*=global_ is 1.21. **(D)** Fitness effect of *AKL1* mutation in all homozygote backgrounds in the suloctidil environment, partitioned by genotypes at *MKT1* (D30G), *RHO5,* and *WHI2.* Initial SSE_*b*=1_ / SSE_*b*=global_ is 1.31. **(E)** Median relative fit ratio between regressions with fixed slope of *b*=1 and *b*=global, as function of number of epistatic terms removed from observed phenotypes. Vertical lines represent interquartile range. Polarity adopted such that *b* ≤ 1. **(F)** Inferred fitness effect of *PMA1* S234C mutation in 4NQO environment across all haploid backgrounds. Epistatic terms interacting with *PMA1* are completely removed from genotype fitnesses, then added back sequentially (from largest to smallest). Bottom-right: full-model (inferred) and observed genotype fitnesses, respectively. Grey line is regression slope. **(G)** Scatterplot and histograms of FCT regression slopes for all data, and number of epistatic terms sufficient to recapitulate them. Horizontal lines in histogram indicate means. Arrows, letters indicate populations presented in previous panels. Polarity adopted such that *b* ≤ 1.

We found that FCTs are common in our landscapes: across all ploidies, environments, and loci, ~44% of regression slopes deviate substantially from 1 (i.e., *b* ≤ 0.9 or *b* ≥ 0.9^-1^; these deviations are all significant; Fig. 3B, see histogram; Fig. S13 and S14). However, FCTs were not universal for fitness-affecting mutations: of the 49 examples across ploidies and environments of mutations with additive effects of magnitude ≥ 0.5%, 18 were associated with 0.9 < *b* < 0.9^-1^ (Fig. 3B).

By partitioning background genotypes by the presence or absence of specific mutations, we can determine whether FCTs are truly “global” (i.e., they transcend these partitions and any corresponding idiosyncratic interactions; Fig. 3A, middle), or are instead fundamentally idiosyncratic (i.e., they emerge from regression across partitions shifted in φ_Mut_ versus φ_WT_ space by sparse interactions with specific background loci; Fig. 3A, right). When we partitioned FCTs by the presence or absence of interacting mutations in the background, we found several instances where the idiosyncratic model clearly explains the fitness-correlated trend. For example, the effect of the G10S mutation in *RHO5* at 37°C exhibits a clear FCT (*b* = 0.76) (Fig. 3C). However, we can partition points by the presence of interacting *WHI2* and *AKL1* alleles in the background. Doing so shows that pairwise interactions with these alleles cause systematic shifts in φ_10S_ vs φ_10G_ space, with each partition assuming a slope near 1. Thus, over a range of background fitnesses, a FCT in the effect of the G10S emerges from these specific idiosyncratic interactions (Fig. 3C, S11). In the case of the homozygous *AKL1* S176P mutation in suloctidil, we observe a similar decomposition of a FCT (*b* = 1.29) when partitioning genotypes according to the presence of three interacting loci in the background (*MKT1, RHO5,* and *WHI2*) (Fig. 3D, S11). However, in other cases it is less clear whether the FCT can be partitioned in this way, and since deeper partitions tend to reduce background fitness variance and limit our confidence in regression slopes, a different approach is required to characterize the extent to which idiosyncratic terms cause FCTs across our data.

To investigate this question, we therefore analyzed the effect of removing specific idiosyncratic epistatic terms on the overall fitness-correlated trends. To do so, for each focal locus (in each ploidy and environment) we first calculated the weighted sum of squared errors (*41*) of observed fitnesses from the global regression line (SSE_*b*=global_) and from a fitted line of slope 1 (SSE_*b*=1_, which corresponds to no FCT). We then set the largest epistatic term to zero and recalculated the expected fitness of each resulting genotype (assuming all other terms and residuals are non-zero), again obtaining both SSE_*b*=global_ and SSE_*b*=1_. If the fitness-correlated trend arose from a global effect, we expect that SSE_*b*=global_ would be less than SSE_*b*=1_ even as terms are removed. Instead, we found that, after removing the effect of just a few terms, a regression with a fixed slope of *b*=1 typically fit the data better than the *b*=global FCT slope (Fig. 3E, S11, with FCT threshold set to *b* ≤ 0.9 or 0.8)), approaching the fit of an unconstrained regression that minimizes SSE (i.e., the final slope approaches 1, Fig. S10). This indicates that the apparent FCT arises from these few idiosyncratic interactions, even for global slopes very different from 1. While we also documented cases where *b*=global fit the data better than *b*=1 even after removing many terms, we expect most if not all these instances may be due to measurement error, since they tend to arise in ploidies and environments where the data is noisier (Fig. S17).

To further evaluate whether idiosyncratic interactions between these mutations are sufficient to generate FCTs, we performed the converse analysis, this time with genotype fitnesses as predicted by our model of additive and idiosyncratic epistatic terms. Instead of removing the effects of epistatic terms one at a time, we first stripped from the model all interactions involving the focal locus, yielding perfectly linear points of slope 1 when plotting φ_Mut_ vs φ_WT_. We then added interactions one by one to our fitness prediction, from largest to smallest, and examined the resulting slopes. As shown in Fig. 3F for the haploid *PMA1* S234C mutation in 4NQO, adding just a handful of terms associated with 3 background loci recapitulates a strong FCT. Repeating this analysis with all our mutations shows that, on average, just 4 idiosyncratic interactions (primarily pairwise) are sufficient to recapitulate the full-model FCTs (a slope within 0.01 of the global slope, Fig. 3G, orange; see SI), which is far lower than the total number of inferred terms (median of 53) but represents on average 89% of the potential variance explained that could have been added (Fig. S12). Thus, while fitness-correlated trends are real and likely have important biological consequences, our data demonstrate that apparent fitness-mediated epistasis can readily emerge from remarkably few low-order idiosyncratic interactions.

Since the landscapes we study here have no natural polarization (i.e., neither allele is the assumed wildtype), we cannot comment directly on why earlier studies of global epistasis have more commonly found negative than positive FCTs (when plotting Δφ versus φ_WT_). However, this distribution of FCT directions is important because it may underly the ubiquitous trend of declining adaptability observed across laboratory evolution experiments (*29*). The observed bias towards negative trends may arise from asymmetries in the average sign of epistatic interactions between mutations away from extant high-fitness genotypes relative to their reversions, which theory has predicted should arise from idiosyncratic interactions (*19*, *20*). In addition, we note that choosing polarizations at random will lead to more negative than positive FCTs across the full parameter space (see extended discussion in the SI).

Regardless of the cause of any asymmetry in the direction of fitness-correlated trends, our results support recent theoretical arguments that fitness-mediated epistasis can emerge as the generic consequence of widespread idiosyncratic interactions, rather than reflecting a global fitness-mediated coupling of mutations. Indeed, at least in our system, we see that fitness-correlated trends can arise even from a relatively small number of low-order interactions. We note that landscapes involving other types of variation (e.g., within a single protein or pathway or along the line of descent in a single lineage (*21*)) may exhibit different patterns, though we may expect these scenarios to involve an even stronger role for idiosyncratic interactions. More generally, we emphasize that idiosyncratic epistasis and global fitness-mediated effects are not mutually exclusive, and while fitness-correlated trends can be explained by the former in our system, in other cases both effects may contribute. However, our results suggest that nonspecific global epistasis may not be the primary driver of patterns of declining adaptability in laboratory evolution experiments, and this has general implications for the ways in which epistasis constrains evolutionary trajectories.

## Acknowledgments

We thank the Bauer Core facility at Harvard, Gautam Reddy, Boris Shraiman, and members of the Desai lab for experimental assistance and comments on the manuscript. We thank Artur Rego-Costa for his help in creating Fig. 2C. Computational work was performed on the Odyssey cluster supported by the Research Computing Group at Harvard University.

## Author contributions

Conceptualization: ANNB, CWB, YS, JIR, MMD

Methodology: ANNB, CWB, YS, JIR

Investigation: CWB, ANNB

Formal analysis: CWB, ANNB

Visualization: CWB, ANNB

Funding acquisition: MMD

Resources: MMD

Supervision: MMD

Writing – original draft: CWB, ANNB, MMD

Writing – review & editing: CWB, ANNB, MMD

## Data and materials availability

Data described in the paper are presented in the supplementary materials. Raw sequencing data will be made publicly available at the NCBI Sequence Read Archive (accession no. TBD), and all analysis code will be made available from Github. Strains used in this study are available upon request.

## Supplementary Materials

Materials and Methods

Supplementary Text

Figs. S1 to S17

Table S1

References (*1–41*)

Data S1 to S3

