## Supplementary material for "Idiosyncratic epistasis leads to global fitness-correlated trends"

### Supplementary Information

References 32

#### List of Tables

S1 Mutations constructed in the experiment . . . . . 4

### 1 Design and construction of the combinatorial CRISPR gene-drive system and library

#### 1.1 Strains

The two parental strains used in this study, YAN548 and YAN564, differ at their mating type and are derived from the BY4742 [1] (S288C: MATa, his3Δ1, ura3Δ0, leu2Δ0, lys2Δ0) with several modifications required for our combinatorial CRISPR gene-drive strategy. We chose to work in this background due to its history in studies of epistasis in yeast [2] and ease of transformation [3].

S288C is a poor sporulator [4], and we introduced the RME1 promoter allele known to increase sporulation efficiency (ins-108A) in BY4742, creating YAN404. YAN407 was generated from YAN404 by mating-type switching using a centromeric plasmid carrying the HO endonuclease (pAN216a_pGAL1-HO_pSTE2-HIS3_pSTE3_LEU2). We then introduced the Cre recombinase under the control of the galactose promoter at the YBR209W locus using Delitto Perfetto [5], yielding YAN525 and YAN526. The CAN1 gene was subsequently replaced with a mating type reporter construct [6] (pSTE2-SpHIS5-pSTE3-LEU2) which expresses the HIS5 gene from *Schizosaccharomyces pombe* (orthologous to the *S. cerevisiae* HIS3) in MATa cells, and the LEU2 gene in MAT$\alpha$ cells.

Cas9 was introduced close to the HO locus under the control of an estradiol-inducible promoter [7] (HO::SpCas9-B112-ER), generating the final strains YAN548 and YAN564. Preliminary work has shown that 2 µM $\beta$-estradiol is sufficient for robust Cas9 induction.

Starting strains containing specific mutations were constructed using dsDNA oligo-mediated repair using Cas9-mediated double-strand break. To do so, we created a centromeric plasmid carrying the URA3 gene that expressed the guide-RNA. Yeast cells were grown with $\beta$-estradiol to induce Cas9, and transformed at log-phase with the guide-RNA expressing plasmid and a double-stranded DNA oligonucleotide with the desired mutation. Cells were then recovered on SD-URA with $\beta$-estradiol to maintain expression of Cas9 and the guide-RNA. A parallel transformation can be done to assess the targetting efficiency as an efficient guide-RNA usually leads to far fewer surviving colonies during the transformation due to the toxicity of unrepaired Cas9-mediated double-strand break. Large colonies from the transformation were then grown in YPD overnight and spread on media containing 5-FOA (1 g/L) to counterselect the plasmid expressing the guide-RNA. All strains were then verified by Sanger sequencing.

#### 1.2 Mutations and their selection

Mutations for our combinatorially-complete fitness landscape were chosen based on several factors. First, we used prior information from published and unpublished experiments that suggested fitness effects for our mutations in at least one environment. Second, due to the need to minimize guide-RNA recognition after the desired mutation is made, we focused on amino acid changes because synonymous mutations could also be incorporated. Third, mutations were chosen that would target a variety of cellular processes to maximize our ability to detect global epistasis. Finally, mutations were chosen that could be efficiently made and not negatively impact our CRISPR-Cas9 system described here (i.e., mutations should not make strains sterile, impair sporulation, or impact galactose metabolism).

| Mutation | Sequence Information | Reference |
|---|---|---|
| WHI2 L262S<br>Chr XV | Guide RNA: ATGGATATGTTGTGCTCCTC<br>L262S DNA: GAcATGagtTGtTCCTCCGGA<br>L262L DNA: GAcATGcTaTGtTCCTCCGGA | [8] |

| | | |
|---|---|---|
| PMA1 S234C<br>Chr VII<br>Essential gene | Guide RNA: TGCTATTACTGGTGAATCTT<br>S234C DNA: ACTGGTGAATgccTtGCTGTC<br>S234S DNA: ACTGGTGAATCccTtGCTGTC | [9] |
| MKT1 D30G<br>Chr XIV | Guide RNA: ATGGTTGACGTCTATATCCA<br>D30G DNA: ACCCTGGgaATtGAtGTtAAC<br>D30D DNA: ACCCTGGAcATtGAtGTtAAC | [10] |
| RHO5 G10S<br>Chr XIV | Guide RNA: ATAATTGGTGATGGTGCAGT<br>G10S DNA: ATatcaGAcGGaGCAGTAGGT<br>G10G DNA: ATaGGaGAcGGaGCAGTAGGT | Our lab |
| AKL1 S176P<br>Chr II | Guide RNA: TCGCGATGGATCAAGGACAC<br>S176P DNA: CCTGTGcCtcTaATtCAcaGa<br>S176S DNA: CCTGTGTCtcTaATtCAcaGa | [11] |
| BUL2 L883F<br>Chr XIII | Guide RNA: CACAAACACGTTTCAAGATT<br>L883F DNA: TGCCCAATtTcGAgACtTGT<br>L883L DNA: TGCCCAATtTgGAgACtTGT | [12] |
| FAS1 G588A<br>Chr XI<br>Essential gene | Guide RNA: AATCGGTAGACCACCTTTAT<br>G588A DNA: ATCGcacGtCCtCCaTTATT<br>G588G DNA: ATCGGacGtCCtCCaTTATT | [13] |
| NCS2 H71L<br>Chr XIV | Guide RNA: CTGAATCAGAATGTGATAAG<br>H71L DNA: CTCCCCTTgagtttgagtGA<br>H71H DNA: CTCCCCTTgagtCAcagtGA | [14] |
| SCH9 P220S<br>Chr VIII | Guide RNA: TCTAATGGTCCTGAGTCACT<br>P220S DNA: AAcGGatCaGAaTCACTAGGC<br>P220P DNA: AAcGGaCCaGAaTCACTAGGC | [15] |
| RPI1 E102D<br>Chr IX | Guide RNA: GTAATGAATGCTATATCCTC<br>E102D DNA: GAGCCTGAcGAcATtGCtTTC<br>E102E DNA: GAGCCTGAaGAcATtGCtTTC | Our lab |

Table S1: Mutations constructed in the experiment. Lower case letters represent mutated sequences with respect to the wild-type DNA.

#### 1.3 Construction of guide crRNA plasmids

Our combinatorial CRISPR gene-drive system allows a hierarchical construction of guide crRNA arrays into a benign locus, by taking advantage of Cre-Lox recombination. Previously, we identified three orthogonal and unidirectional recombination sites that are necessary for our design. Briefly, our gene-drive system makes use of three types of recombining plasmids with three distinct pairs of drug markers, which we refer to as type HygMX-KanMX, KanMX-NatMX, and NatMX-HygMX. The three drug markers - HygMX, KanMX, and NatMX - are resistance cassettes for hygromycin B, G418, and nourseothricin, respectively, and differ additionally by the use of paralogous TEF promoters and synthetic terminators as in [16]. Each type is based on an HO-targeting plasmid pAN3H0a (Figure S1), which contains the two drug marker cassettes for selection as well as homologous sequences that lead to integration of insert sequences with high efficiency. The insert sequences between the two drug cassettes contain one of 10 guide-RNA cassettes (each with an SNR52 promoter mutated at non-functional regions to reduce the rate of unintended homologous recombination, the guide-RNA, the structural RNA element and the SUP4 terminator [17]). In

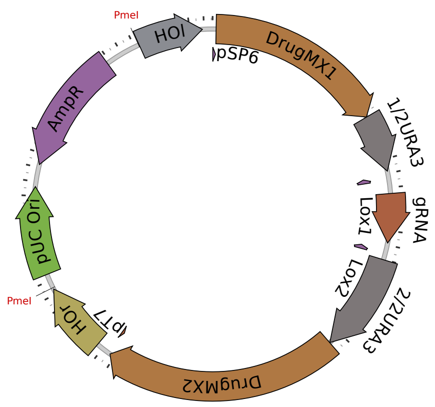

Figure S1: gRNA integration plasmids. We use three integration plasmids with different drug markers, Lox sites, and URA3 frameshift configurations as explained in Section 1.3.

addition, each drug marker is linked to their own half of URA3 (frameshifted for each drug such that the first half of URA3 only functions properly when the correctly framed second half of URA3 is used) which contains a splice donor or acceptor (from QCR10 [18]) and their own orthogonal Lox site (LoxP, Lox2272, or Lox5171, with arm mutations to allow only a single recombination event between them [16]). In the configuration found at integration, the URA3 is not functional. However, when recombined properly by Cre recombinase, a configuration which brings like drug markers on the same chromosome (HygMX-HygMX, for example) will produce a functional URA3, which we can select with media lacking uracil and counterselect with media containing 5-FOA.

This system allows diploids created by mating two strains with compatible marker configurations to be selected on media containing all three drugs (described later in section Section 1.5). Compatible configurations will always include a common drug that will yield a functional URA3 after recombination. For example, the HygMX-KanMX configuration is compatible with KanMX-NatMX (which will form HygMX-NatMX and KanMX-KanMX after recombination) or with NatMX-HygMX (which will form NatMX-KanMX and HygMX-HygMX after recombination). The recombined 'landing pads' are thus compatible with each other (for example, HygMX-NatMX is compatible with NatMX-KanMX, which when recombined will form HygMX-KanMX and NatMX-NatMX).

#### 1.4 Final barcoding procedure

To allow bulk phenotyping of the strains, we introduced a 22mer DNA barcode (16 random nucleotides and 6 known spacer nucleotides) alongside a complete LYS2 ORF at the *LYS2* locus via homologous recombination in the AKL1-RPI1 double-mutant strains prior to the final mating step.

To produce a library of uniquely barcoded plasmids, we generated an entry vector with 702 bp homologous region upstream of the LYS2 deletion, the deleted 293 bp region immediately upstream of the ORF, the 4179 bp LYS2 ORF, and then a 39 bp tGuo1 synthetic terminator. Downstream of this terminator was a primer-binding site, pBC1, followed by the ccdB gene, which is toxic in *E. coli* strain DH10B. This gene was followed by 300 bp of semi-random DNA sequence (used as "filler" for obtaining PCR bands distinct from primer dimer bands), the pBC2 primer-binding site, and 589bp homologous to the region immediately downstream of the LYS2 deletion. Barcodes

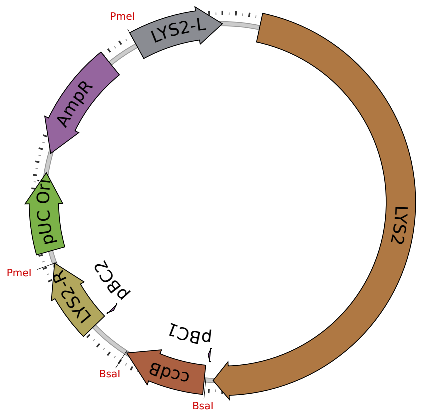

Figure S2: Barcoding plasmid before barcode insertion. We integrate a random barcode at the LYS2 locus to uniquely tag each individual in the pool.

were cloned into this plasmid at the ccdB locus via Golden Gate assembly [19, 20] in 8 independent replicates, separately cloning in DH10B via electroporation and selecting on LB+Ampicillin sodium salt (100 µg/mL) agar plates (1% tryptone, 0.5% yeast extract, 0.5% sodium chloride, 1.5% agar) after an hour of recovery in SOC media (2% tryptone, 0.5% yeast extract, 8.56mM sodium chloride, 2.5mM potassium chloride, 10mM magnesium chloride, 10mM magnesium sulfate, 20mM glucose). Plates, which bore at least 30,000-40,000 transformant colonies each, were each scraped and cultured in 5mL LB+Amp media prior to miniprepping to isolate plasmid.

To barcode AKL1-RPI1 double mutants, we first isolated 10 individual colonies of each of the 4 possible double-mutant genotypes. We split these 10 colonies into two sets of 5. Each set of 5 colonies was cultured, pooled, and transformed with one of the eight barcode plasmid libraries, which had previously been cut with PmeI to linearize the region for integration. Transformants were selected on SD-Lys agar plates and, to the best of our abilities, individually picked into SD-Lys media for continued purifying growth.

#### 1.5 Hierarchical mating procedure

The basic procedure for a cycle of mating, drive, recombination, and sporulation is as follows:

Strains with compatible guide-RNA "landing pads" and opposite mating type were mixed to generate diploids in YPD plus ampicillin (100 µg/mL) via mating for 12-24 hours. Cells were then passaged to YPG (1% yeast extract, 2% peptone, 2% galactose) plus ampicillin liquid media containing hygromycin B (at 300 µg/mL), geneticin (at 200 µg/mL), and nourseothricin (at 100 µg/mL) to select for diploids, with selection sustained for at least 3 generations. Cells were then transferred to YPG containing all four drugs and at least 2 µM $\beta$-Estradiol to induce Cre-recombinase and Cas9, with selection for at least 10 generations. This generates homozygous diploids at the loci targetted by Cas9, and combines the guide-RNA from the homologous HO loci onto the same chromosome. The cells were then grown in SD-Ura with $\beta$-estradiol for at least 15 generations to select for successful Cre-Lox recombinants. They then were induced to sporulate by 16-24 h growth in YPA (1% yeast extract, 2% peptone, 1% potassium acetate) followed by culture in SPO media (1% potassium acetate, 0.005% zinc acetate). After 3-5 days of sporulation, haploids containing all the mutated loci and recombined gRNA loci were selected with at least 15 generations of growth in S/MSG-D (1.67% yeast nitrogen base lacking ammonium sulfate, 1% monosodium glutamate, 2% dextrose) lacking histidine or leucine (selecting for MATa and MAT$\alpha$ respectively), containing two of the three antibiotic drugs (depending on the landing pad configuration) and 1g/L 5-FOA to counterselect diploids. Finally, selected populations were screened for "leakers" by growing a single colony or a small number of cells (less than about 1000) in YPD, followed by a transfer into YPD containing the drug to which the desired haploids should not be resistant. Only specimens sensitive to this third drug were preserved as a frozen archive and then passaged into the next mating-drive-recombination-sporulation step.

In practice, this procedure included a variety of manipulations. This range of manipulations demonstrates that our method is flexible and can be adapted to work within various technical constraints. For example, when handling few strains, microtiter plates are not necessary and the protocol can be performed in standard culture tubes. In the case of the initial double-mutant mating, for instance, mating was in most cases conducted on YPD-agar patches, which were then scraped and transferred into the YPG diploid selection media. All other matings were conducted in about 90 µL YPD liquid media in wells of 96-well round-bottom microtiter plates. Similarly, selection of haploids after sporulation was sometimes conducted in microtiter plates (128 µL total volume), and other times by streaking to individual colonies on SD-Leu or SD-His agar plates (without 5-FOA counterselection). For all cycles except the experiment's final cycle, individual colonies were isolated and screened at the conclusion. Finally, depending on the scale of the cycle, diploid selection, recombinant selection, presporulation, and sporulation steps were conducted in either microtiter plates (shallow for selections (128 µL media), 2-mL deep-well plates for presporulation and sporulation) or test tubes (5 mL media unless otherwise stated).

Presporulation: Microtiter plate-based presporulation was carried out by pipetting 20 µL saturated SD-Ura culture into 480 µL of YPA. Plates were shaken at 1050 rpm at 30°C for 24 hours under a breathable membrane (VWR, 60941-086) before sporulation. Tube-based presporulation was carried out by inoculating 5 mL YPA with 150 µL saturated SD-Ura culture and incubating on a roller drum at 30°C for 16-24 hours.

Sporulation: Microtiter plate-based sporulation was carried out by pelleting presporulated cells at 2000 g for 2 min, washing by resuspension in 400 µL water, pelleting once again, and resuspending in 400 µL sporulation media. These plates were sealed with a breathable membrane, secured with tape to plate shakers, and shaken at 1350 rpm at room temperature for 4-5 days. Tube-based sporulation was carried out by pelleting tube-presporulated cell cultures and resuspending in 2 mL sporulation media, incubating at room temperature on a roller drum for 3-4 days.

Homozygotes from the final cycle were incubated for 5 generations in YPD+Amp prior to archival freezing, but only after fully selecting for recombination of the landing pad loci with SD-Ura+$\beta$-estradiol.

The final 20 generations of haploid selection in the final cycle were conducted in typical haploid selection media, but lacking lysine, in order to select only for those haploids which retained the barcode next to the LYS2 marker (which segregated in a Mendelian fashion).

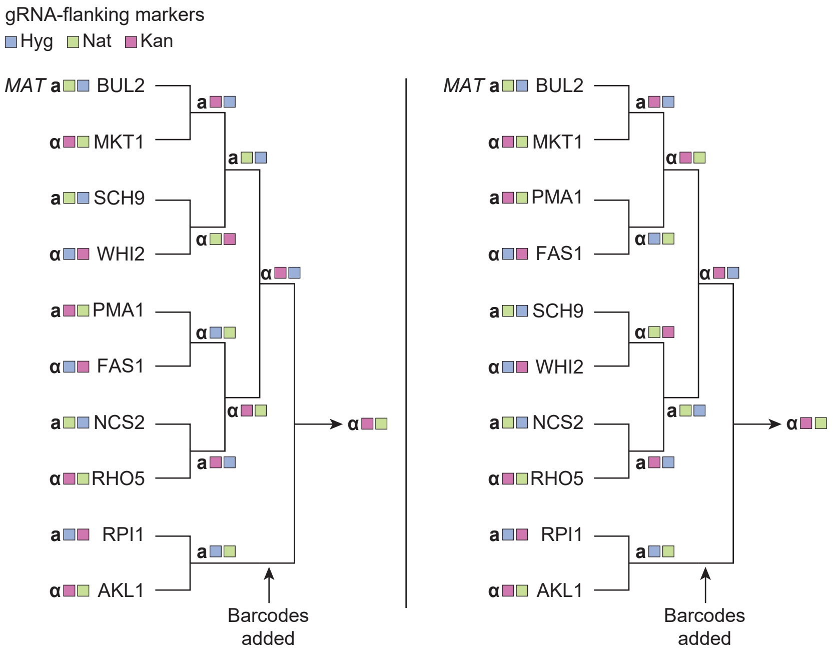

Figure S3: Parallel mating scheme. Biological replicates of the final strains were created via different mating paths.

### 2 Genotype verification

#### 2.1 Whole-genome sequencing

To verify the lack of systematic off-target Cas9-mediated modifications, and to rule out pervasive aneuploidies, we performed whole-genome sequencing on 96 random clones (3 random wells from each of 32 plates which contained 64 different strains based on the mating procedure outlined in Section 1.5) [21] and sequenced each to approximately 100x coverage. This identified a single case of aneuploidy with elevated read counts at three chromosomes that were consistent with disomy. In addition, it identified five credible non-synonymous mutations occurring on more than 1 strain (strongly indicating that the mutations were introduced in the hierarchical mating scheme described in Section 1.5). Two of these (in ERG6 and QRI7) were present in just two strains each, and the other three (in SPT7, HSL7, and FRS1) were present in 5, 6, and 33 strains, respectively. In addition, some extra mutations were identified in single clones, which is not inconsistent given the rate of mutations during meiosis (70% of clones had no mutations, 10% had one, 5% had two, and the rest had poor sequencing coverage leading to what we believe are bad variant calling). These results suggest that Cas9 does not introduce a gross excess of off-target mutations in the genome, and that although unintended mutations do occur in our system (due to Cas9 or meiosis) these mutations are unlikely to dominate the estimation of parameters for modeling the fitness landscape. Notably, as explained in Main Text, biological replicates (independent crosses) were typically in agreement with each other.

To understand whether the mutations in SPT7, HSL7, and FRS1 may have systematic effects on our genotypes, we looked at whether they were present exclusively on any single- or double-mutant backgrounds. We found that the FRS1 mutation was present across most backgrounds, but the mutations in SPT7 and HSL7 were only present on specific AKL1-RPI1 backgrounds. Notably, fewer than half of the instances of these backgrounds in our WGS data bore these mutations.

#### 2.2 Locus-specific multiplex PCR

To genotype the entire haploid library at all 10 primary loci and 3 putative segregating off-target loci (FRS1, SPT7, and HSL7), we pursued a multiplexed strategy. We began by lysing all 2048 wells (not all of which contained cells) with 20 µL yeast lysis buffer (5mg/mL Zymolyase 20T, 100 mM Sodium Phosphate pH 7.4, 10 mM DTT) and 5 µL of cells straight from the freezer stock. The enzymatic reaction was placed at 37°C for at least 45 min and then at 95°C for 2 min. The released DNA could then be stored in the freezer overnight.

Immediately prior to the first-round PCR, we boiled these products for a minute to mix the lysates. We then added and mixed in 25 µL of water to the lysis products to dilute and facilitate liquid handling. Then, we added 2 µL of this lysis product to the PCR master mix for the first round PCR, mixing after addition. This master mix was for a 25 µL Phire reaction and contained 1.3 µL of pooled 100 µM primers. These primers represented all 13 loci. The 13 primers that added N7 adapters to the amplicons were common across all wells. The 13 primers that added S5 adapters contained 6 bp inline indices. These indices existed in 8 versions, each unique to a different set of 4 plates in the library (54°C annealing, 45s extension). These primers may be found in Data Table S1.

The following day, PCR round 1 products were combined into 4 pools, taking 4 µL from each well. We cleaned up these pools with a 1x bead purification step (AMPure beads by Beckman Coulter) (starting volume = 42 µL , eluted in 35 µL ). We used KAPA polymerase for a second round of 25 µL PCRs to anneal unique pairs of S5/N7 indices to the amplicons across 4 reaction plates, using 2 µL of purified round 1 product (63°C annealing, 45s extension). Several unsuccessful reactions were repeated as necessary with diluted template.

Round 2 reaction products were then pooled and cleaned via gel extraction, followed by a final bead purification step to remove any remaining small fragments.

The library was sequenced on a NextSeq mid output lane resulting in an average coverage of about 2700x per locus per well in the genotype library. Loci varied in their overall coverage: the average coverage per BUL2 locus was just about 80x, whereas the average coverage for WHI2 was about 7300x. Other than BUL2, all other loci had an average coverage of at least 400x.

Some loci for specific wells were missing from our dataset, or otherwise had very low coverage. To patch these holes in our genotyping data, we amplified with locus-appropriate primers in a first-round reaction to anneal S5/N7 adapters. This reaction used Phire polymerase (54°C annealing, 45s extension) and 2 µL of diluted lysate as template. These reaction products were cleaned up with 1x Ampure beads and eluted in 30 µL water. We took 2 µL of this reaction product into the second round KAPA Hifi PCR reaction, which annealed pairs of S5 and N7 indices unique to each reaction (63°C annealing, 45s extension). Each reaction product was then cleaned up separately using 0.8x Ampure beads on 6 µL of reaction product diluted in 10 µL water. The final product was eluted in 25 µL and pooled for sequencing on a MiSeq Nano lane.

#### 2.3 Counting alleles for each locus in each well

Once we received the Illumina reads, we counted the number of reads of each allele at each locus in each well. To do this, we followed a procedure similar to [22], examining each read in each 8-well sequencing library (corresponding to individual fastq files) in turn. First, we checked that the first 6 bp of read 1 corresponded to a 6-bp inline index, allowing for 1 bp of mismatch. Then, we evaluated

read quality by ensuring that the quality score of the 22bp downstream from the inline index was at least 25. If a read met these conditions, we identified the locus associated with the read by checking for the presence of a characteristic 8-bp sequence either upstream or downstream of the defined allele, allowing identical matches only. For reads matching an identifiable locus, we extracted the 20- to 23-bp allele, sequentially using a list of decreasingly stringent regular expressions (using the python regex module [23]):
'(left 8bp)(length of allele)(right 8bp)',
'(left 8bp)(length of allele-2,length of allele+2)(right 8bp)',
'(left 8bp){e≤1}(length of allele)(right 8bp){e≤1}',
'(left 8bp){e≤1}(length of allele-2,length of allele+2)(right 8bp){e≤1}',
For lists of the exact alleles and 8-bp sequences searched, see Data Table S1.

Overall, fewer than 0.5% of reads were excluded on the basis of these criteria, with no more than 1.2% for a single library.

All alleles that occurred at least 10 times in at least one well AND were present at at least 1% frequency for the corresponding locus in at least one well were given a unique identifier and assigned as either a WT, Mut, or Other allele. "WT" alleles included properly repaired pseudo-WT alleles plus other versions with some or even none of the desired synonymous changes. This includes loci in which it appears no gene drive occurred (i.e., sequences identical to the unmutated parental BY sequence). "Mut" alleles included any with the desired missense change, regardless of the presence or absence of other synonymous alleles. "Other" alleles included those whose amino acid sequence matched neither the WT nor Mut sequence, including errant missense changes and frameshifts. Any remaining alleles were grouped together and designated "na."

#### 2.4 Statistical inference of gene-drive failures

One difficulty of verifying locus correctness by PCR in the final haploid library is that the strains are not clonal (they are derived from the Cas9 gene-drive hierarchical mating procedure, see Section 1.5). Thus, we needed to remove wells that had evidence of a mixture of genotypes, or strong evidence of the incorrect genotype. We noticed that our multiplex PCR verification protocol in Section 2.2 produced evidence of genotype mixtures at a higher rate than anticipated. However, we observed that these supposedly incorrect wells were found more frequently when post-first-round PCR pools were "mixed" at a given locus (i.e., were expected to have both WT and Mut alleles present). This indicated to us that primers from the first-round PCRs were leaking through, thus incorrectly indexing the reads, and/or PCR chimeras were forming.

We developed a statistical model to accurately estimate the true mixture proportion within each well. For each post-first-round pool, we calculated the pool-wide frequencies of all alleles in that pool (based on their unique identifiers). Then, we modeled a pool-wide probability $p$ that a given read is a "true" read and not a chimeric read. We assume that the expected frequency of a false allele in a given well will be $(1-p)\cdot$ the poolwide frequency of that allele, whereas a true allele in a given well will have an expected frequency of $p+(1-p)\cdot$ the poolwide frequency of that allele. For a range of values of $p$, constraining $p$ to be at least 50%, we calculated the probability of the data under a multinomial model and obtained the maximum likelihood estimate of $p$. As necessary, we constrained the likelihood surface to satisfy the constraint that all alleles must be present at a frequency between 0 and 1, inclusive.

After obtaining these adjusted allele frequencies, we set out to distinguish which wells were acceptably versus unacceptably "pure." Since rates of apparent chimera formation varied significantly across loci, we developed a separate purity threshold for each locus. We did this by sorting wells by the percent of non-dominant alleles at a given locus, excluding "na" alleles. We plotted these proportions against the ascending rank on the x axis, forming a "hockey stick"-like curve that shoots upward at the high end of the distribution. We found the "elbow" of this curve, i.e., the proportion of non-dominant allele at which the curve is furthest from a "hypotenuse" line connecting the end of the handle to the tip of the blade of the proverbial hockey stick. We obtained the following thresholds from this approach:

Approximate thresholds

BUL2 3.04%, gives 93.3% pure

FAS1 1.20%, gives 95.1% pure

MKT1 1.13%, gives 96.6% pure

NCS2 1.94%, gives 96.3% pure

PMA1 2.29%, gives 91.8% pure

RHO5 1.58%, gives 94.8% pure

SCH9 0.64%, gives 94.5% pure

WHI2 0.45%, gives 92.9% pure

AKL1 3.27%, gives 95.3% pure

RPI1 2.36%, gives 95.1% pure

For the sake of comparison, we note that overall drive failure rates inferred from sequencing the quadruple and octuple mutants – which was not done in a pooled, chimera-genic way – were close to 2%. In addition, many gene drive events that in fact failed may not be counted here, since a failed drive event that yields the unmutated WT allele when the WT allele is desired will be retained.

All told, 1282 wells matched their expected genotype at all loci (67.8%). Since we had biological replicates of each genotype, 875 out of 1024 possible genotypes (85.4%) were represented among these wells. See Data Table S2 for a complete list of wells, barcodes, and genotypes that passed these filters.

#### 2.5 Other genotyping

In addition to genotyping the final products of the experiment, we genotyped one or more mutated clones per genotype after each cycle of mating, drive, recombination, and selection before proceeding with the next cycle.

Genotyping of the double mutants (after cycle 1) was conducted via Sanger sequencing and visual examination of traces for the expected alleles.

Genotyping of the quadruple mutants (after cycle 2) and octuple mutants (after cycle 3) was conducted via next generation sequencing.

Quadruple mutants were lysed in 50 µL yeast lysis buffer (5 mg/mL Zymolyase 20T (Nacalai Tesque), 1 M sorbitol, 100 mM sodium phosphate pH 7.4, and 20 mM DTT), boiled at 95°C for 2 minutes and 2 µL lysed cells were taken into a 25 µL Phire polymerase PCR reaction with 1.25 µL each of the 4 pairs of appropriate primers for 4 loci, respectively (54°C annealing, 30s extension). After this first round of PCR, we purified the product with 0.8x beads and did a second round KAPA Hifi polymerase PCR (25 µL ) to append unique S5, N7 indices to each colony isolate. The final product was purified with 0.8x beads once again and sequenced libraries on MiSeq Nano 2x150bp.

Octuple mutants were lysed with 5 µL of saturated culture in 50 µL yeast lysis buffer as previously described. The boiled lysis product was diluted two-fold, 2 µL of the lysis was used in into 24 µL Phire polymerase PCR reaction containing 1 µL of each of 16 10 µM primers, each of which add the S5, N7 adapter sequences (54°C annealing, 30s extension). Round 1 PCR products were purified via bead cleanup at 0.8x beads ratio, and eluted with 25 µL water. Before cleanup, some of wells were diluted with an additional 10 µL to bring volume up (evaporation of PCR reactions is frequent), then 12 µL taken as starting volume for cleanup. We then performed the round 2 PCR reaction with unique pairs of S5 and N7 primers for each well, taking 2 µL of cleaned up DNA template into a 25 µL KAPA reaction (63°C annealing, 45s extension time). Round 2 PCR products were diluted with an additional 10 µL water and pooled (3-4 µL of each well). We then performed a 0.7x bead cleanup and submitted the final purified pool for NextSeq Mid throughput 1x150bp lane.

### 3 Combinatorial indexing and sequencing of barcodes

#### 3.1 Combinatorial pooling and sequencing

To map the barcodes to individual wells, we took a combinatorial indexing approach. Uniquely barcoded AKL1-RPI1 double mutants were cultured in the central 64 wells of 32 96-well microtiter plates (rows A-H, columns 3-10). With the help of a Biomek liquid handler, we took 10 µL of each well-mixed well culture into either of two new 96-well plates, in which wells had been seeded with 30 µL of YPD to facilitate automated liquid dispensing. 70 µL of pooled culture from each well of these two plates was used to form 8 row-specific pools, and the process was repeated form 8 column pools. Each pool contained approximately 1.1 mL of culture. Separately, for each of the 32 plates, 20 µL from each of 64 wells was pooled to form 32 plate pools of about 1.3 mL each.

To prepare libraries for sequencing, we extracted genomic DNA from each of the 48 pools, eluting in 50 µL water. In an initial PCR step using primers 5xx>pBC1-F and 7xx>pBC2-R, we amplified the barcode loci in each pool, attaching S5 and N7 adapters to each amplicon. For these reactions, we used 0.5-5 µL of genomic DNA in a 25 µL Q5 reaction (34 cycles, 54°C annealing, 45s extension). After purifying amplicons via a cleanup with 0.8x ampure beads and eluting into 33 µL water, we performed a second round of PCR with 1 µL of purified DNA template and unique pairs of S5 and N7 primers (KAPA 50 µL reaction, 34 cycles, 63°C annealing, 45s extension). Final PCR products were pooled, with 2 µL of each plate pool and 8 µL of each row and column pool (total volume about 200 µL ). Half of this was taken for a 2-sided bead selection, first with 0.5x beads, and next with 0.2x more beads for a 0.7x selection.

Libraries were sequenced on a NextSeq mid-output lane yielding an average coverage of about 8700 reads per barcode per pool.

#### 3.2 Barcode assignment to single wells

Combinatorial indexing allows one to uniquely triangulate a barcode to a specific well. However, errors due to sequencing, apparent cross-contamination due to chimeric reads or lower read coverage for some particular combinatorial pool can make some assignments ambiguous. We therefore performed this assignment using a greedy procedure. First, barcodes that uniquely map to a single well were identified. This yielded 2332 barcodes (out of 2348) that mapped to 2029 wells. Evidently, some wells contained multiple barcodes that stem from imprecise colony picking. 16 barcodes appeared to map ambiguously to multiple wells. Manual inspection found that 12 of these could be explained by spurious reads in other pools, which meant we only had to remove four wells with conflicting barcodes.

We additionally found about 40 wells that appeared to grow extremely slowly in SD-Ura+$\beta$-estradiol+Amp, perhaps due to picking petite colonies. All were of the same AKL1-RPI1 genotype (AKL1 176S, RPI1 102D) and from the same barcode transformant pool, leading us to believe this may be due to private mutations in one of the 5 replicate pooled colonies. We manually identified, removed, and repicked these remaining wells from the opposite transformant pool of that same AKL1-RPI1 genotype, in which we had seen no issues. In addition, we repicked about a dozen barcode transformant colonies for wells with unassigned barcodes.

The second set of barcodes was assigned again by combinatorial indexing, this time with only 8 rows and 8 columns, and some spurious remaining wells that did not have a well-defined barcode were also confirmed by Sanger sequencing.

### 4 Bulk phenotyping

#### 4.1 Growth experiments

The complete frozen pool was grown in 5 mL YPD by inoculating approximately $10^7$ total cells to produce the starting population. We then diluted these populations by 1:$2^7$ daily by passaging 781 µL into 5 mL fresh media (of some particular environment) in 15 mL culture tubes on roller drums. Whole population pellets, obtained from 1.5 mL of saturated culture, were stored immediately at -20°C for later sequencing. As previously described [16], this protocol results in about 7 generations per day, with a daily bottleneck size of about $10^8$ in most assay environments. We performed two replicates of each assay and sampled for 49 generations (7 timepoints). Only 5 timepoints (representing 7, 14, 28, 42, and 49 generations) were sequenced.

The six environments chosen were: YPD + 0.4% acetic acid (YPDA), YPD + 6 mM guanidium chloride (gu), YPD + 35 µM suloctidil (suloc), YPD + 0.8 M NaCl (salt), YPD at 37°C (37C), and SD + 10 ng/mL 4NQO (4NQO). (All environments besides 37C were at 30°C.) The YPDA environment was chosen because preliminary experiments suggested that it had a tendency to reveal phenotypic variance and it previously had been studied in our lab ([16]). Gu was chosen because of its known large target size from separate work in our lab which identified a change in sign for the effect of the MKT1 D30G mutation. Suloc and 4NQO were chosen because previous work in our lab showed these environments to have low genotype correlation with other YPD-based environments. 37C and salt were chosen because several of the genes under study were previously reported to be mutated in evolution under that stressor or be a QTL in that stressor (NCS2 in high temperature; PMA1, RPI1, and RHO5 were all mutated in NaCl evolution experiments; see Table 1.2).

The degree of the stressor in suloc, YPDA, salt, and 4NQO environments was chosen empirically to maximize the stress while still permitting 7 generations of growth per day over the entire phenotyping assay.

#### 4.2 Amplicon barcode sequencing

Genomic DNA from cell pellets were processed as in [16]. Briefly, DNA was obtained by zymolyase-mediated cell lysis (5 mg/mL Zymolyase 20T (Nacalai Tesque), 1 M sorbitol, 100 mM sodium phosphate pH 7.4, 10 mM EDTA, 0.5% 3-(N,N-Dimethylmyristylammonio)propanesulfonate (Sigma, T7763), 200 µg/mL RNAse A, and 20 mM DTT) and binding on silica mini-preparative columns with guanidine thiocyanate buffer (4 volumes of 100 mM MES pH 5, 4.125 M guanidine thiocyanate, 25% isopropanol, and 10 mM EDTA). After binding, the columns were washed with a first wash buffer (10% guanidine thiocyanate, 25% isopropanol, 10 mM EDTA) and then a second wash buffer (80% ethanol, 10 mM Tris pH 8), followed by elution into elution buffer (10 mM Tris pH 8.5). 1.5 mL of pelleted cells eluted into 100 µL routinely provided about 1-2 µg of total DNA.

PCR of the barcodes was performed using a two-stage procedure previously described to attach unique molecular identifiers (UMIs) to PCR fragments (see [16] for a detailed protocol). Primers used in the first-stage PCR contained a priming sequence, a 7-12-nucleotide multiplexing index,

8 random nucleotides as UMIs, and an overhang that matched the Tn5 transposome. These two primers had the configurations P1 = TCGTCG GCAGCG TCAGAT GTGTAT AAGAGA CAGNNN NNNNNY YYYYYY AAGGTA CGATTC TGACGC A, P2 = GTCTCG TGGGCT CGGAGA TGTGTA TAAGAG ACAGNN NNNNNN YYYYYY YAGTTG TCTCTG CTCTCG CTA. Here N corresponds to degenerate bases used as UMIs, and Y corresponds to multiplexing indexes.

These primers anneal on either side of the barcode sequence integrated just downstream of LYS2, at the pBC1 and pBC2 sites, respectively. After attachment of molecular identifiers to template molecules during three PCR cycles (20 µL Q5 Polymerase reaction, 50°C annealing, 30s extension), the first-stage amplicons were cleaned using Ampure beads using an automated liquid handling protocol established for a Biomek FXp, with 1.25x Ampure beads, eluting in 35 µL . Of the elution of this clean-up, 30 µL was used directly as template for the second-stage PCR with primers that contained multiplexing indexes and adapters that anneal to the Illumina flowcells (P5 and P7 primers). After 35 PCR cycles (50 µL KAPA Hifi Polymerase reaction, 63°C annealing, 30s extension), these final products were then purified using Ampure beads, quantified, and pooled to approximately equimolar concentration. The PCR products were sequenced with a NovaSeq S1 full flow cell (Illumina) by paired-end sequencing (2 x 50 bp, reading 80 bp from the P1 direction and 20 bp from the P2 direction).

We first processed our raw sequencing reads to identify and extract the indices and barcode sequences as in [16]. Using the barcodes previously identified in Section 3.2, we can make "corrections" to reads with sequencing errors by direct lookup of the lowest Levenshtein distance to the dictionary of verified barcodes.

Finally, we can calculate the counts of each error-corrected true barcode by removing duplicate reads, using the unique molecular identifiers from the first-stage PCRs. Frequencies calculated from these counts are used to infer fitnesses for all segregants, as explained in Section 4.3. After all filtering, our final mean sequencing coverages were over 1500 reads per barcode per timepoint per replicate (averaged across all assays).

#### 4.3 Fitness inference for time-dependent barcode frequencies

Strain fitnesses can be inferred from relative barcode frequencies over time (see Refs. [16] and [24] for expanded information on joint inference of fitnesses using barcode frequencies). Briefly, fitnesses are regressed as the change in relative log frequencies of strains against a selected reference per generation. This parameter is approximately the difference in instantaneous growth rate between lineages under exponential growth. Most genotypes in our data are represented by more than one barcode in the same assay (representing biological replicates), and each barcode was measured in two technical replicates. In theory, we could jointly infer the biological replicates and constrain their fitnesses to be equal. This would yield, for a combinatorially complete landscape, exactly $2^N$ fitnesses which could be fit exactly with $2^N$ coefficients (later described in Section 5.1). However, strains with the same desired genotype may not always be identical at all other loci in the genome (due to new mutations or off-target effects). By only performing the joint inference on technical replicates, variance left unexplained by a full model containing $2^N$ coefficients can be regarded as biological variation at other loci and some measurement error (described in more detail in Section 5.2). This joint inference is intuitively similar to a weighted average of the two technical measurements, with weights proportional to the evidence within each replicate (which is a combination of the number of reads and the number of timepoints measured). A standard error for the inferred fitness parameter can be obtained through the inference process by the square root of the inverse of the Fisher information at the maximum likelihood. This standard error can be interpreted as the error that can be attributed to the (overdispersed) binomial sampling error. For our analyses, we removed datapoints with standard error above 1.

##### 4.4 Comparison between technical, biological replicates

As shown in Figure S3, biological replicates were made for all final strains in the experiment by proceeding through a parallel mating scheme. However, due to gene-drive failures, some strains were not found in replicate, and it may be useful to ask the following questions: 1) How trustworthy are the strains without replicates? and 2) What is the average effect of unintended mutations introduced within our cross? To answer these questions, we can compare the inferred fitnesses of technical (comparing the same barcode across assays) and biological replicates (comparing barcodes that correspond to the same expected genotype).

Decomposing the observed phenotypic variance due to measurement error can be done by the standard reliability estimates. The Pearson's correlation coefficient between two technical replicates is an estimate of the $R^2$ between the true fitness value and one fitness measurement for the barcode. If one takes the mean of the r technical replicates, then:

$$\frac{\sigma_{err}^2/r}{\sigma_{gen}^2 + \sigma_{err}^2/r} = \frac{1 - \langle \rho_{r_i,r_j} \rangle}{1 + (r-1)\langle \rho_{r_i,r_j} \rangle}. \qquad (1)$$

Decomposing the phenotypic variance due to extra variance in the genetic component can be done by a similar process, by comparing the measurement values between strains bearing different barcodes but expected to have the same genotype. Here, to perform this calculation, we constrain ourselves to pairs of strains with the same genotype that each have a single barcode in their well so that a single comparison can be made. The correlation coefficient between biological replicates can be interpreted in a similar way to technical replicates, but the deviation from 1 here will reflect both error due to extra variation in the genome and error due to measurement error (but without tube-to-tube variation). For the purpose of our manuscript, we assume that this tube-to-tube variation is negligible.

In plots of technical and biological replicates, density-based coloration was determined by calculating each point's mean distance to its five nearest neighbors. Distances were transformed using the scikit MinMaxScaler() function and plotted with normalized colors based on a reversed viridis colormap.

Technical and biological replicate comparisons for all data can be viewed in Fig. S4.

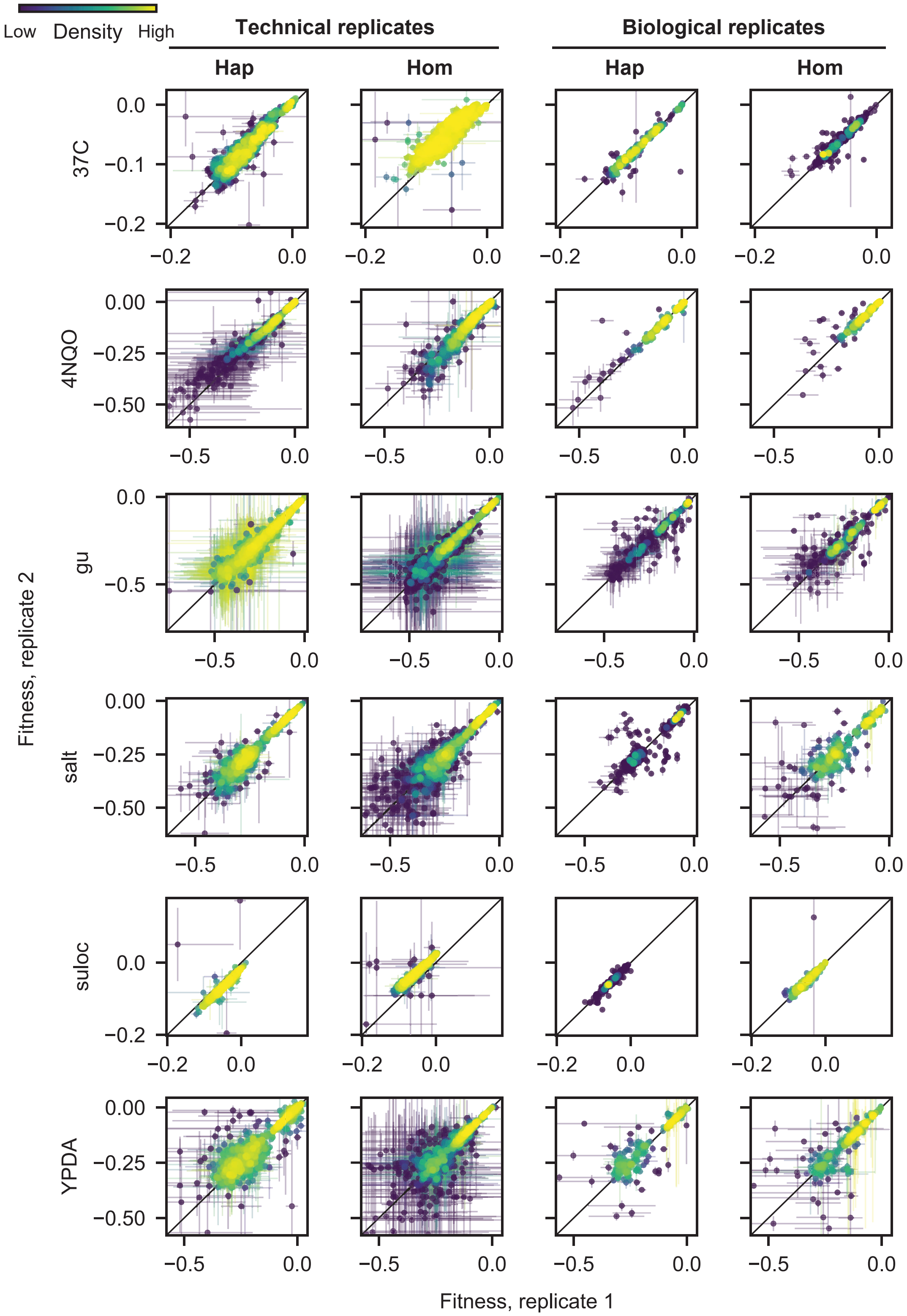

.

Figure S4: Comparison of technical and biological replicates for all ploidies and assay environments

### 5 Quantitative analysis of epistasis

#### 5.1 Estimation of parameters

We model fitness $\phi$ as a function of the the underlying genotype which can be expressed as a sum of combinations of N biallelic loci $x_1$, $x_2$,..., $x_N$ that take on values $x_i = \pm 1$.

$$\phi = \overline{\phi} + \sum_i s_i x_i + \sum_{i>j} s_{ij} x_i x_j + \sum_{i>j>k} s_{ijk} x_i x_j x_k + ... \tag{2}$$

This modeling framework casts additive first-order terms as the background-average effect of the mutation, which is distinct from the effect of the mutation on some arbitrary wild-type genotype. The terms $s$ represent half the fitness difference between groups of individuals with and without the mutation, or alternatively the expected deviation from the mean, positive or negative, for groups with or without the mutation respectively. Pairwise epistatic effects are the background-average perturbation that can be fit beyond the additive first-order term, and higher order epistatic terms are similarly modeled. This view offers several advantages: 1) if one decides to choose a particular genotype as the "wild-type", only the signs of the terms need to change; 2) each coefficient is estimated by partitioning half of the genotypes (each coefficient corresponds to a distinct slice of the data), meaning each coefficient is equally powered; and 3) the coefficients are in principle orthogonal from each other (there is no expected collinearity between the genotypic values of any pair or combination of coefficients). This means that there is no "order" of coefficient fitting (one does not have to fit the additive terms first), and fitting one coefficient does not influence another.

Coefficients from the equation above can always be estimated by least-squares regression when all $2^N$ genotypes have a phenotypic measurement, though we note that we have in practice on average more than 1 phenotypic measurement per genotypes due to our biological replicates. However, we may expect this formula to be sparser: not all mutations should have an effect, and not all pairs of mutations should have a pairwise epistatic effect. We can regularize the estimation procedure to yield a sparse subset using the LASSO procedure, which penalizes the least-squares regression by the sum of absolute magnitudes of coefficients:

$$\min_s \left\{ ||\phi - \hat{\phi}||_2^2 + \lambda ||s||_1 \right\}. \tag{3}$$

In the absence of collinearity (as stated above, our formulation has no collinearity between parameters), the LASSO operation is known to be consistent and asymptotically selects the correct subset of non-zero parameters [25]. Sparsity is controlled by the $\lambda$ parameter, which can be found by cross-validation (in our case, 5-fold cross-validation was performed to reduce the extent of overfitting). This approach removes coefficients that are approximately the same scale as the noise. To provide 95% confidence intervals on the LASSO estimates, we performed 500 bootstrap resampling with replacement of the data followed by model selection.

As discussed previously in Section 2.1, we identified extra mutations present in multiple strains (FRS1, SPT7, HSL7). Because the SPT7 and HSL7 mutations likely occurred during the mating process (Section 1.5), they may lead to specific signals of epistasis if they themselves have an effect. We briefly assessed this possibility by plotting the distribution of fitnesses for individuals with and without the mutation (constraining on the backgrounds in which the mutations were identified). In visually examining these plots, we were unable to find evidence of a systematic effect for these mutations. Therefore, these mutations were removed from consideration before building the model by LASSO. On the other hand, FRS1 was likely present in one of the original parents of the experiment and thus was found in approximately 50% of final strains. Though we did identify a possible effect for this mutation in some environments, because it is not systematically distributed across the library, it is only expected to affect one of the higher order epistatic terms. (We cannot distinguish the effect of the epistatic term for the combination of strains that have FRS1 mutated and the effect of the FRS1 mutation). However, note that we have produced strains in replicate. Thus, the effect of the FRS1 mutation is unlikely to be consistently found in the same strains, and its signal will therefore be unlikely to dominate the epistatic term.

In general, the broad-sense heritability captured by the model is very high as both biological and technical replicates show high correlation (see Fig. S4). Thus, correlation of fitness measurements between environments can reveal the similarities between model coefficients. If measurement noise was too great such that it would dilute the correlation coefficients, then comparison between the predicted fitnesses may provide a better picture of environmental similarities (given that the coefficients were adequately estimated).

##### 5.2 Variance partitioning

The phenotypic variability in the dataset can be partitioned into various components to quantify their relative importance. In our experiment, we are interested in not just the broad-sense heritability due to our focal loci ($H^2$, or the variability due to genetic components), but also in the heritability due to specific additive and epistatic components. When the model coefficients are orthogonal, the phenotypic variance due to genetic components is trivially obtained by the sum of squares of each coefficient:

$$\sigma^2_{\text{gen}} = \sum_i s_i^2 + \sum_{i>j} s_{ij}^2 + \sum_{i>j>k} s_{ijk}^2 + \ldots \tag{4}$$

Partitioning the variance by subsets of coefficients – for example partitioning by first order terms or pairwise epistatic terms – is therefore straightforward.

$$\sigma^2_{\text{1st}} = \frac{\sum_i s_i^2}{\sigma^2_{\text{gen}}} \tag{5}$$

$$\sigma^2_{\text{2nd}} = \frac{\sum_{i>j} s_{ij}^2}{\sigma^2_{\text{gen}}} \tag{6}$$

However, we note that the coefficients are estimated from the data, and variance partitioning in this manner produces a bias. Removal of this bias is the major motivation behind mixed linear models that estimate narrow-sense heritability [26]. This caveat is not a major concern for our study, though, since extra sources of variation are either negligible (all the phenotypes are measured simultaneously in the same tube) or can be well estimated (measurement error can be estimated by replication). None of these extra sources of variation are expected to fundamentally alter only some of the coefficients or some subset of coefficients, and thus these relative partitions are expected to be unbiased.

#### 6 Analysis of fitness-correlated trends

##### 6.1 Fitting regression slopes to determine fitness-correlated trends

Fitness-correlated trends (FCTs), such as diminishing returns or increasing costs, have often been analyzed by regressing the fitness effect of a mutation, $s = \Delta\phi = \phi_{mut} - \phi_{wt}$, against the fitness of the background in which it occurs, $\phi_{wt}$. We refer to this as the $\Delta\phi$ formulation: we say that there is no FCT if $\Delta\phi$ is constant over a wide range of background fitness, while a negative relationship between $\Delta\phi$ and $\phi_{wt}$ corresponds to diminishing returns/increasing costs (and a positive relationship corresponds to increasing returns/diminishing costs). However, care must be taken when performing this analysis, because when we regress $\Delta\phi$ against $\phi_{wt}$, measurement errors in $\phi_{wt}$ will lead to a negative correlation even in the absence of true fitness-correlated trends [27].

A further complication with this formulation is that the regression slope we obtain depends in a complex way on the polarization we choose for the mutation (i.e., which allele is considered the wild-type and which is the mutant). To see this, consider the following simple linear model for $\Delta\phi$ as a function of $\phi_{wt}$:

$$\Delta\phi \equiv \phi_{mut} - \phi_{wt} = a_1 + b_1\phi_{wt}, \tag{7}$$

and the analogous model for the fitness effect of the reversion, $\Delta\tilde{\phi}$, as a function of $\phi_{mut}$:

$$\Delta\tilde{\phi} \equiv \phi_{wt} - \phi_{mut} = a_2 + b_2\phi_{mut}. \tag{8}$$

Fitting data to these models using standard methods for ordinary least-squares, we find that the relationship between the regression slopes $b_1$ and $b_2$ is given by

$$b_2 = -\frac{b_1 + V}{1 + 2b_1 + V}, \tag{9}$$

where we have defined

$$V = \frac{\mathrm{Var}[\Delta\phi]}{\mathrm{Var}[\phi_{wt}]}. \tag{10}$$

We can use these equations to gain some intuition for the effect of $V$ on the regression slopes and their reversions (i.e., a change in polarization). First, $V \geq 0$ by construction, and $V = 0$ only if there is no measurement error or no idiosyncratic epistasis, which in some extreme cases could be interpreted as measurement error for all measurements. As expected, it is only possible to lack an FCT in both polarizations ($b_1 = b_2 = 0$) if $V = 0$. Of note, the numerator of $V$ can be decomposed to $\mathrm{Var}[\phi_{mut}] + \mathrm{Var}[\phi_{wt}] - 2Cov(\phi_{mut}, \phi_{wt})$, which shows that without a specific relationship between fitnesses of individuals with and without the mutation, $V > 0$, and an FCT will always emerge in at least one of the two polarizations.

Since in practice $V$ is always positive, we can see that, as shown in Figure S5 and from Equation 9, when $b_1 \geq 0$, then $b_2 < 0$, no matter the difference in scale of $V$ and $b_1$. Thus, in practice, increasing returns (or diminishing cost) epistasis or no FCT in one polarization of which allele is the "WT" always shows as diminishing returns (or increasing cost) in the reversion (when the allele is considered to be the "Mut" instead).

When $b_1 < 0$, or when there is diminishing returns in this polarization, then the behavior of $b_2$ depends on the scale of $b_1$ and $V$. First, some scenarios lead to $b_2 = 0$, or no FCT in the reversion, and these scenarios occur at the critical boundary where $V = -b_1$. Another critical boundary occurs where $V = -1 - 2b_1$, which leads to an asymptotic boundary where $b_2 \to \pm\infty$. When $b_1 < 0$, only a small region between the critical boundaries leads to $b_2 > 0$ (the reversion is increasing returns or diminishing cost epistasis). Outside the critical boundaries, $b_2 < 0$ and therefore diminishing returns or increasing costs is found in both polarizations of the allele. Thus, across the full space of possible parameters, diminishing returns and increasing costs – both of which present as a negative regression slope – are more likely to emerge than positive regression slopes in this $\Delta\phi$ formulation (though we note that biology may not explore this entire parameter space uniformly), and slopes when mutations are reverted cannot always be anticipated intuitively. We can also ask when $b_2 = b_1$: this will happen when $b_1 = -0.5V$. Because $V \geq 0$, this will only happen when $b_1 < 0$ (and therefore $b_2 < 0$). Another fact from this equality is that if $b_2 = b_1$,

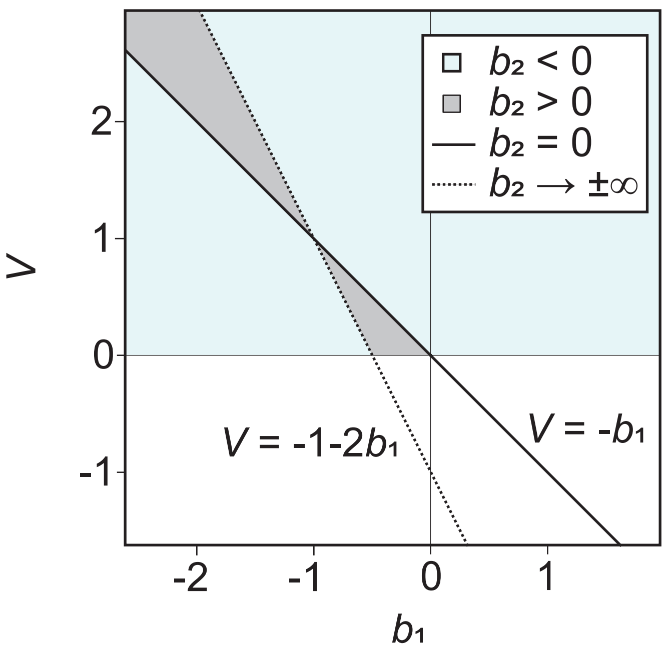

Figure S5: Relevant regimes for slopes and their reversions in the $\Delta\phi$ formulation. $V$ is from Equation 10, $b_1$ and $b_2$ are least-squares regression slopes when an allele is labeled as the WT allele or the mutated allele (i.e., the reversion).

then the denominator of $V$ has to be equal in the reversion. This means that $b_2 = b_1$ implies $\mathrm{Var}(\phi_{wt}) = \mathrm{Var}(\phi_{mut})$.

Note, these complications are still present when using other regression techniques such as total least squares that take into account measurement errors in $\phi_{wt}$ and $\phi_{mut}$ [28].

In contrast, we can resolve some of these complications by making two changes to the analysis: (1) plotting $\phi_{mut}$ directly against $\phi_{wt}$, and (2) regressing a linear relationship based on the total least squares. Firstly, this approach avoids some problems with correlation in measurement errors. In this formulation (i.e., the $\phi_{wt}/\phi_{mut}$ formulation), measurement errors in both strains (or errors in the dependent and independent variable) are taken into account [28] (we use the standard errors estimated from Section 4.3), and we have the model functions:

$$\phi_{mut} = a_3 + b_3\phi_{wt} \tag{11}$$

and the reversion:

$$\phi_{wt} = a_4 + b_4\phi_{mut} \tag{12}$$

Secondly, this framing and regression method (taking into account errors in both axes) also behaves far more intuitively: the slope in one direction is always the reciprocal of the other (i.e., $b_3 = 1/b_4$).

To obtain some intuition of how to interpret FCTs in this $\phi_{wt}/\phi_{mut}$ formulation, we can first attempt to interpret $b_3 = 1 = b_4$. This only occurs if $\mathrm{Var}(\phi_{wt}) = \mathrm{Var}(\phi_{mut})$, a property of the regression method. As described earlier, this is the regime where $b_1 = b_2 \leq 0$, and $b_1 = b_2 = 0$ only if $\mathrm{Var}(\phi_{mut} - \phi_{wt}) = 0$. Thus, a caveat of this $\phi_{wt}/\phi_{mut}$ formulation is that a slope of 1 does

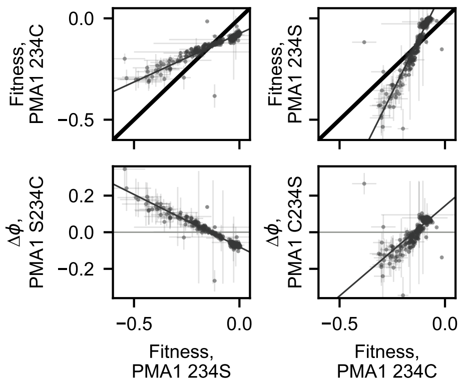

.

Figure S6: Comparison of fitness correlated trends for a simple case where the reversion of the focal mutation is straightforward. Haploid, 4NQO environment.

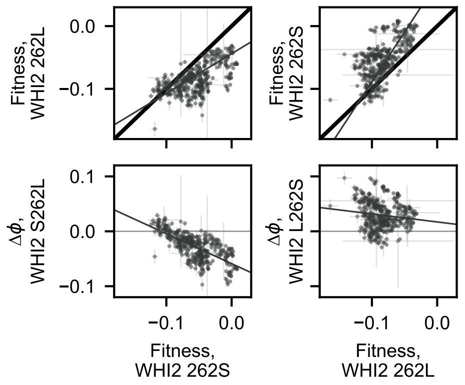

.

Figure S7: Comparison of fitness correlated trends for a complicated case where the reversion of the focal mutation is not intuitive. Haploid, high-temperature environment.

not always indicate the absence of an FCT. In contrast, when $b_3 \neq 1$, then either $b_1 \neq 0$ or $b_2 \neq 0$. This can be shown by the fact that $b_3 \neq 1$ only when $\text{Var}(\phi_{wt}) \neq \text{Var}(\phi_{mut})$. This case necessarily implies $\text{Var}(\phi_{mut} - \phi_{wt}) \neq 0$, which is the necessary condition for $V > 0$.

We summarize these behaviors with some example figures from our data. First, we show an example of intuitive behavior, comparing both regressions and with mutational reversions (Figure S6). In this simple example, regression of the fitness effect of the PMA1 234C mutation leads to a case of diminishing returns and increasing cost epistasis. When the mutation is "reverted," or we regress the effect of the 234S mutation, we obtain the opposite FCT (diminishing costs, or increasing returns). These trends are also well-captured in the $\phi_{wt}/\phi_{mut}$ formulation.

On the other hand, many examples are far less intuitive (Figure S7). In this example, regressing the effect of the WHI2 262L mutation leads to diminishing returns. However, regressing the effect of the reversion (262S) also leads to diminishing returns. In the $\phi_{wt}/\phi_{mut}$ formulation, slopes behave as expected (the reversion is the reciprocal).

Examples where FCTs can only be interpreted in one of the mutational orientation are also found (Figure S8). In this example, the PMA1 234C mutation apparently shows no FCT, while its

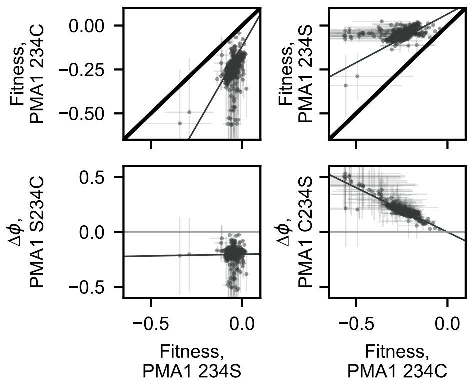

.

Figure S8: Comparison of fitness correlated trends for a case where reversion may be interpreted as having no FCT. Haploid, acetic acid environment.

reversion displays increasing cost epistasis. On the other hand, the $\phi_{wt}/\phi_{mut}$ formulation robustly shows a slope different from 1 and again behaves as the reciprocal when the mutation is reverted. Thus, because different slopes in this formulation do not readily yield an interpretation of the type of FCT (diminishing returns vs increasing returns), we refrain from using these plots for this purpose. Instead, we focus on this formulation's ability to robustly identify FCTs when it exists.

For our analysis, a final complication emerges from having biological replicates. In the parameter estimation above (Section 5.1), this does not pose a problem (there is simply unexplained variation). However, for the purpose of analyzing fitness correlated trends, if strains have two replicates for the wild-type and two replicates for the mutant, then there are 4 possible comparisons and it is no longer clear how to regress this effect of the mutation. To resolve this, we perform the analysis on the average fitness of each genotype, which can be interpreted as the best estimate of the true fitness of the genotype. The standard error of the average genotype fitness was computed as the mean of the errors associated with the fitnesses that were averaged.

We have shown that, in general, if slopes different from 1 are obtained in the $\phi_{wt}/\phi_{mut}$ formulation, then we can interpret the data as displaying FCTs. However, what yields slopes different from 1? If these formulations are readily interchangeable, then we may expect a single idiosyncratic epistatic term involving the focal mutation, positive or negative, to be sufficient. However, we find that this is not the case: in this formulation, we find that this epistatic interaction must also involve a mutation with a non-zero additive effect.

To illustrate this, we begin with a simple schematic considering two loci A and B on top of a background of other mutations with some fitness variance (Figure S9). We denote alternative alleles at these loci as their letter case (A/a, and B/b), and the deviation from the mean fitness between genotypes of alternative alleles for locus A as: $s_A = \phi_A - (\phi_A + \phi_a)/2$. When $s_A = 0$, $s_B = 0$, and $s_{AB} = 0$, then plotting $\phi_A$ vs $\phi_a$ must yield a general "cloud" of points with a slope of 1 (Figure S9, top left panel). Partitioning the cloud of points by genotypes with the $B$ and $b$ alleles, respectively, only yields two superimposed clouds (because the effect of having the mutation at locus B, $s_B$, is zero). When $s_A = 0$, $s_B \neq 0$, and $s_{AB} = 0$, then the two clouds separate themselves along the 1:1 line (Figure S9, top right panel). The regression slope for $\phi_A$ vs $\phi_a$ is still 1. The case where $s_A = 0$, $s_B = 0$, and $s_{AB} \neq 0$ is more complicated. Setting $s_{AB} = E$, a constant, we find the mean deviation in fitnesses $\phi_{AB} = E$, $\phi_{aB} = -E$, $\phi_{Ab} = -E$, and $\phi_{ab} = E$. If we focus on plotting $\phi_{ab}$ against $\phi_{Ab}$, we find that the negative deviation due to the epistatic coefficient for

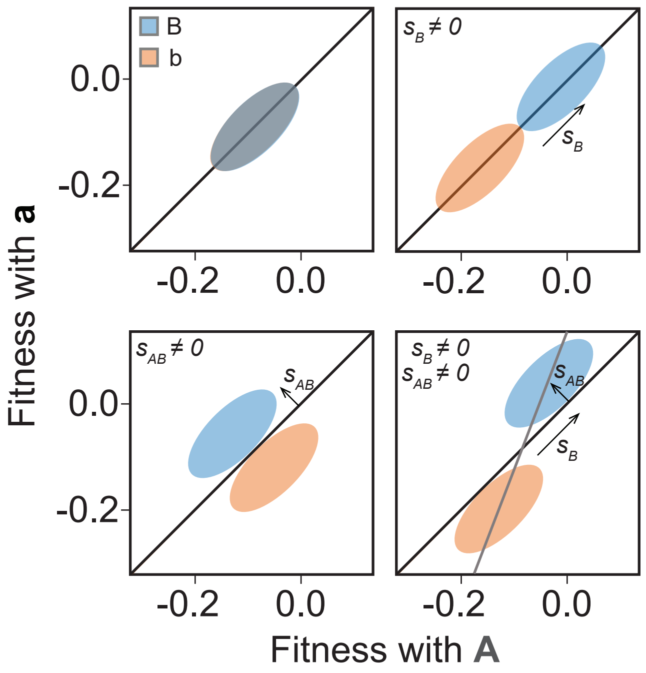

Figure S9: Effect of parameters on global regression in the $\phi_{wt}/\phi_{mut}$ formulation. Clouds shown are for when $s_A = 0$. Top right shows the effect of $s_B \neq 0$, bottom left shows the effect of $s_{AB} \neq 0$ and bottom right shows the effect of both $s_b \neq 0$ and $s_{AB} \neq 0$.

$\phi_{Ab}$ moves the cloud to the left, while the positive deviation due to the epistatic coefficient for $\phi_{ab}$ moves the cloud up. These coordinated movements yield a diagonal movement orthogonal to the 1:1 line. The same logic can be applied to plotting $\phi_{aB}$ against $\phi_{AB}$, however in this case the cloud moves to the right and down. Thus, the two clouds separate themselves in the direction of a slope of -1 when an epistatic term is present (Figure S9, bottom left panel). The regression slope for $\phi_A$ vs $\phi_a$ is still 1 even in this case and will eventually flip to be -1 as clouds separate themselves farther and farther. Putting these orthogonal movements together, we find that the non-zero terms for $s_A = 0$, $s_B \neq 0$, and $s_{AB} \neq 0$ lead to joint cloud movements (Figure S9, bottom right panel). The regression slope for $\phi_A$ vs $\phi_a$ in this final case will never be one. Because these conditions include the sufficient condition for FCTs in the $\Delta\phi$ formulation, our analyses on FCTs with this $\phi_{wt}/\phi_{mut}$ formulation are conservative, and we use this formulation for its advantages: 1) errors in fitness measurements are taken into account for both $\phi_{wt}$ and $\phi_{mut}$, 2) the slope for the mutation reversion is the reciprocal, and 3) slopes different from 1 are always FCTs.

#### 6.2 Decomposition of fitness-correlated trends

To understand whether idiosyncratic interactions lead to fitness-correlated trends, we proceeded down two analytical avenues.

In the first, we examined the observed genotype fitnesses and removed epistatic terms one at a time to see whether slopes converged to 1. Operationally, this involved first finding the global linear regression line that fit the data best for a given locus in a given ploidy and environment. We compared that regression to the best-fit line with slope of 1 by looking at the weighted sum of

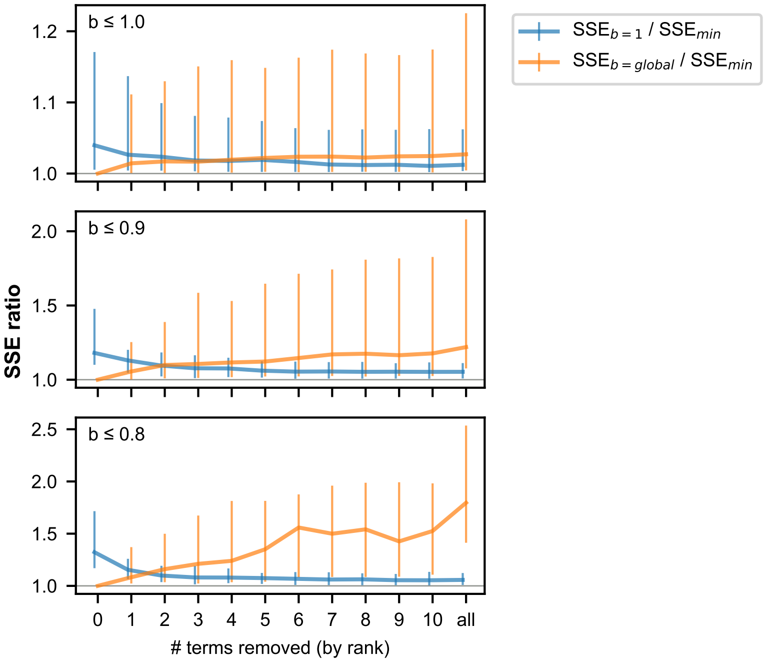

.

Figure S10: Three panels, one for each $b$ threshold, showing the change in the ratios of the SSE for lines of slope $b$ =1 and $b$ =global as compared with the SSE for an unconstrained regression that minimizes SSE. Vertical bars indicate interquartile ranges.

squared errors (SSE), where a lower sum indicates a better fit to the data. After doing this, we found the residual difference between the observed genotype fitnesses and the genotype fitnesses as predicted by our full model of additive and epistatic terms. Then, we set the largest epistatic term involving the focal locus to zero, regenerated the model fitness values, and added the residual differences. To this dataset, we fit a line with the original global slope and a line with the slope 1, again finding the SSE for each. We also fit a totally new regression line that minimized the SSE. We then iterated this process, consecutively removing 10 epistatic terms and re-evaluating the fit of the $b$ =1 and $b$ =global lines each time. Main text Figure 3E shows how the relative fit of these two lines changes across ploidies, environments, and loci. Figure S10 shows how the SSE for $b$ =1 and $b$ =global compare to the minimized SSE as terms are progressively removed, revealing that a slope of 1 tends to approach an idealized fit as terms are removed, while the global slope tends to drift away. Figure S11 provides a more detailed look at how the ratio of SSEs for $b$ =1 and $b$ =global change as terms are removed for each locus in each ploidy and environment.

In a converse analysis, we examined genotype fitnesses generated by our model of additive and epistatic terms. For a focal locus in a given ploidy and environment, we first stripped away all epistatic terms related to interactions between the focal locus and other loci, such that only additive terms and interactions among background loci contributed to the modeled genotype fitnesses. This produced a perfectly straight line with a slope of 1 and an intercept proportional to the background-averaged additive effect of the focal mutation (as described in Section 5.1, this is twice the estimated parameter $s_i$ where $i$ is the focal locus). We ranked the epistatic terms involving the focal mutation by their effect size. Then, starting with the largest, we incorporated one term at a time into the modeled genotype fitnesses. After each term was added, we replotted the fitness of genotypes with

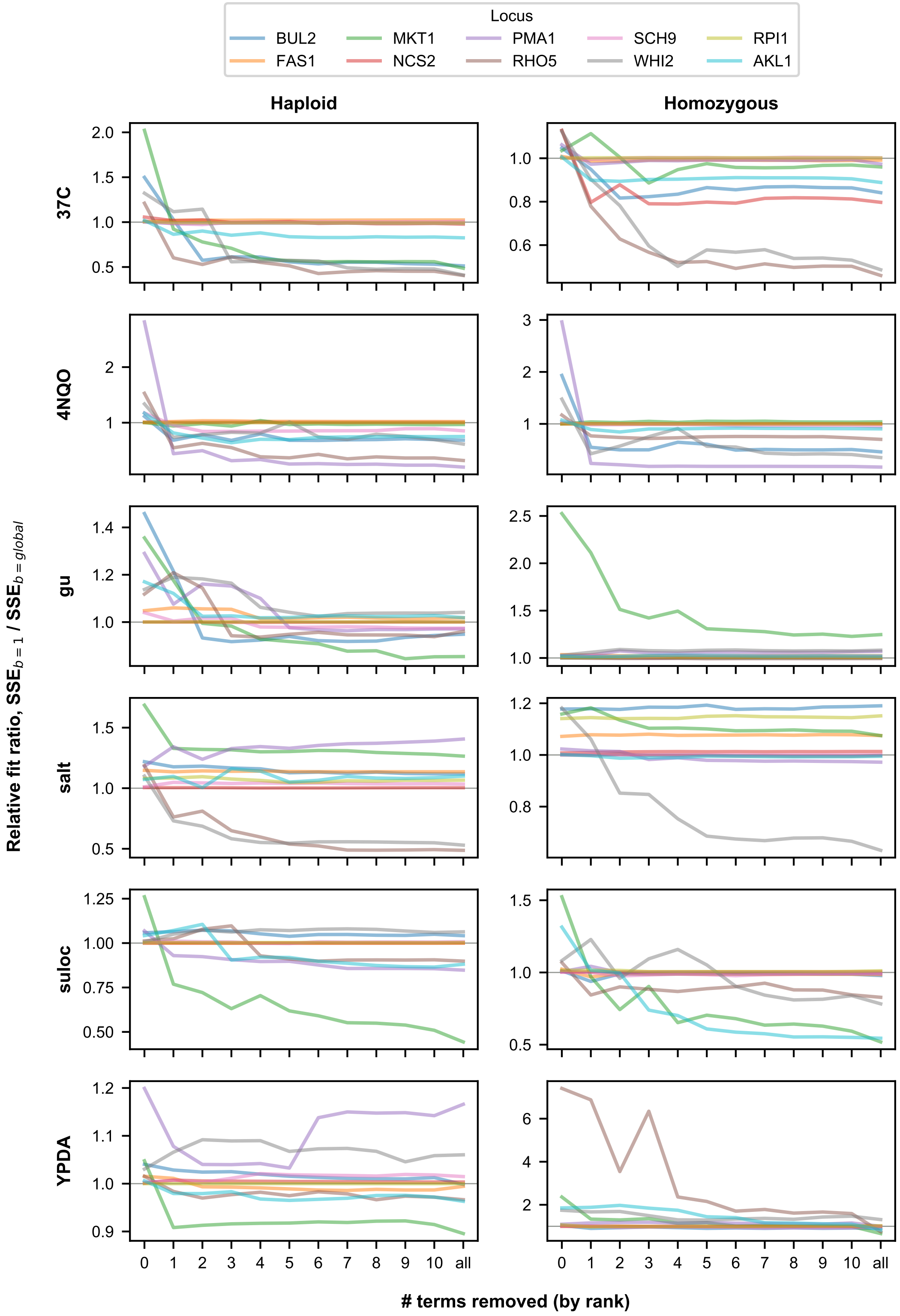

.

Figure S11: One panel for each ploidy and assay environment showing the change in the ratio of the SSEs for lines of slope $b = 1$ and $b =$global for each locus.

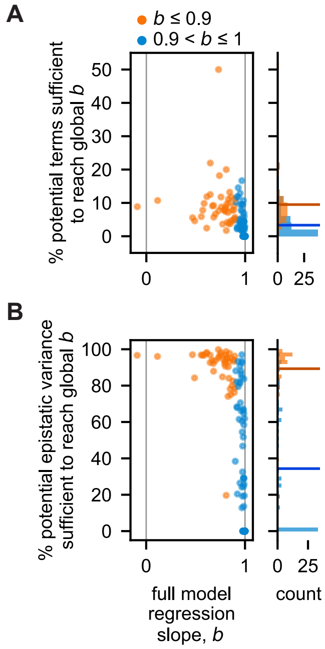

Figure S12: Scatterplot and histograms of regression slopes of FCTs for all data and the percentage of inferred epistatic (A) terms and (B) variance needed to recapitulate them. Horizontal colored lines in the histogram illustrate the mean.

and without a mutation at the focal locus and computed the regression slope.

We defined the number of terms sufficient to recapitulate the observed FCT as the number of added terms required to reach regression slope convergence within 0.01. More specifically, after adding each term, we asked whether the new regression slope differed from each of the previous three regression slopes by less than 0.01. If so, the number of terms required to reach that "plateau" was considered the number of terms sufficient to recapitulate the observed FCT. In a minority of cases, the final "plateau" slope differs from the full-model slope by greater than 0.01, but only in 5 instances by greater than 0.02. Figure S12 presents the fraction of potential epistatic terms and potential epistatic variance sufficient to reach this plateau.

Note that, to permit more consistent comparisons, all loci were analyzed in the mutational direction that placed their regression slopes between -1 and 1. In other words, if plotting genotype fitness with $A$ on the x axis and genotype fitness with $a$ on the y axis gave a slope greater than 1, we would flip the axes such that the slopes would be equal to the reciprocal of the original slope (between 0 and 1).

Plots of $\phi_{wt}/\phi_{mut}$ for all loci can be found in Figure S13 and Figure S14.

#### 6.3 Quantifying the effect of landscape size in the analysis of fitness-correlated trends

The size of the fitness landscape we consider has two important effects on our ability to analyze the origins of fitness-correlated trends. First, as the number of mutations involved increases, the number of potential epistatic interactions between them increases exponentially. This creates more opportunities for idiosyncratic interactions to exist and to produce apparent fitness-correlated trends. We note that this is an average effect: if we happened to choose precisely the set of mutations that

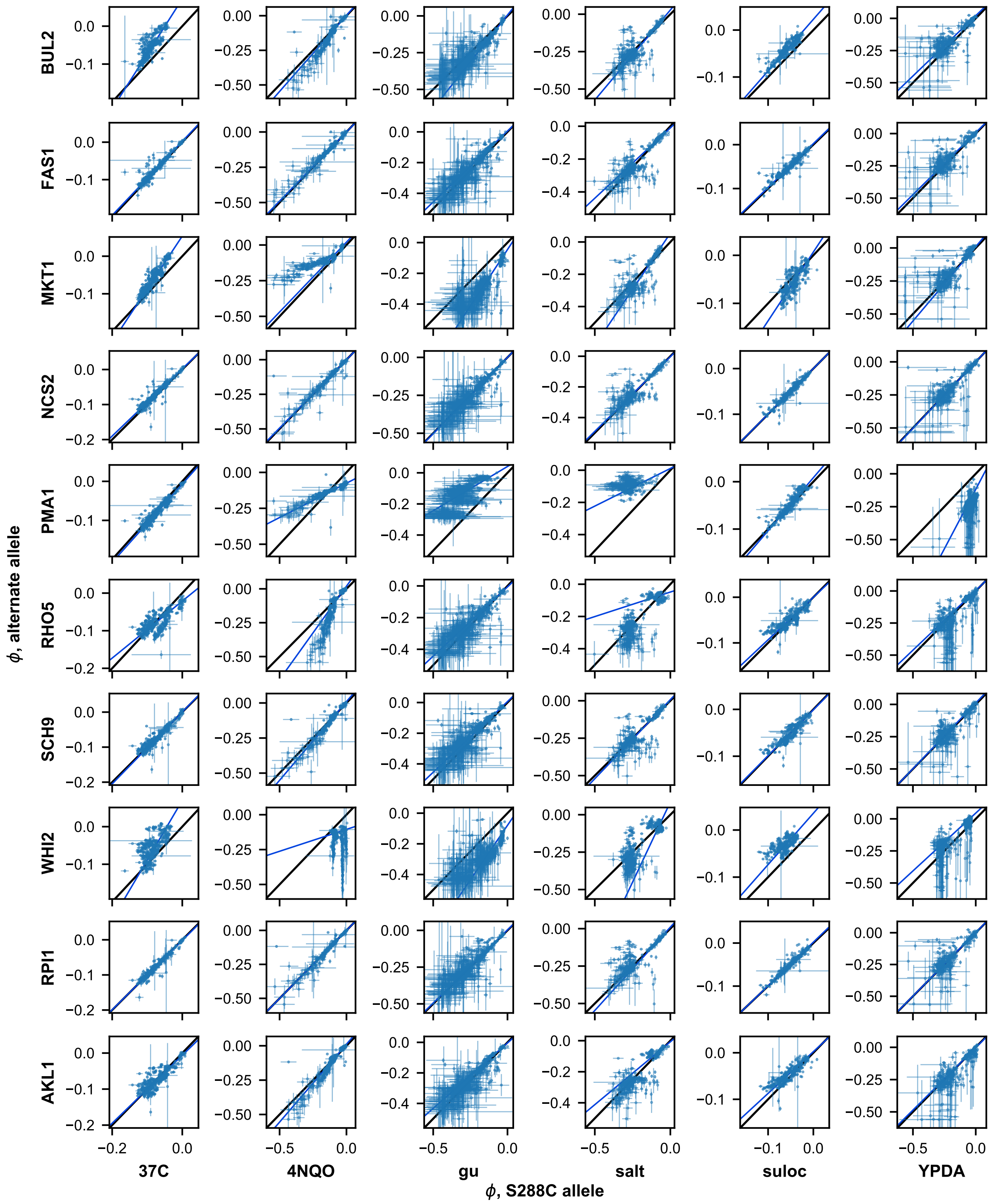

.

Figure S13: Scatterplots of $\phi_{wt}/\phi_{mut}$ for all loci in haploid form.

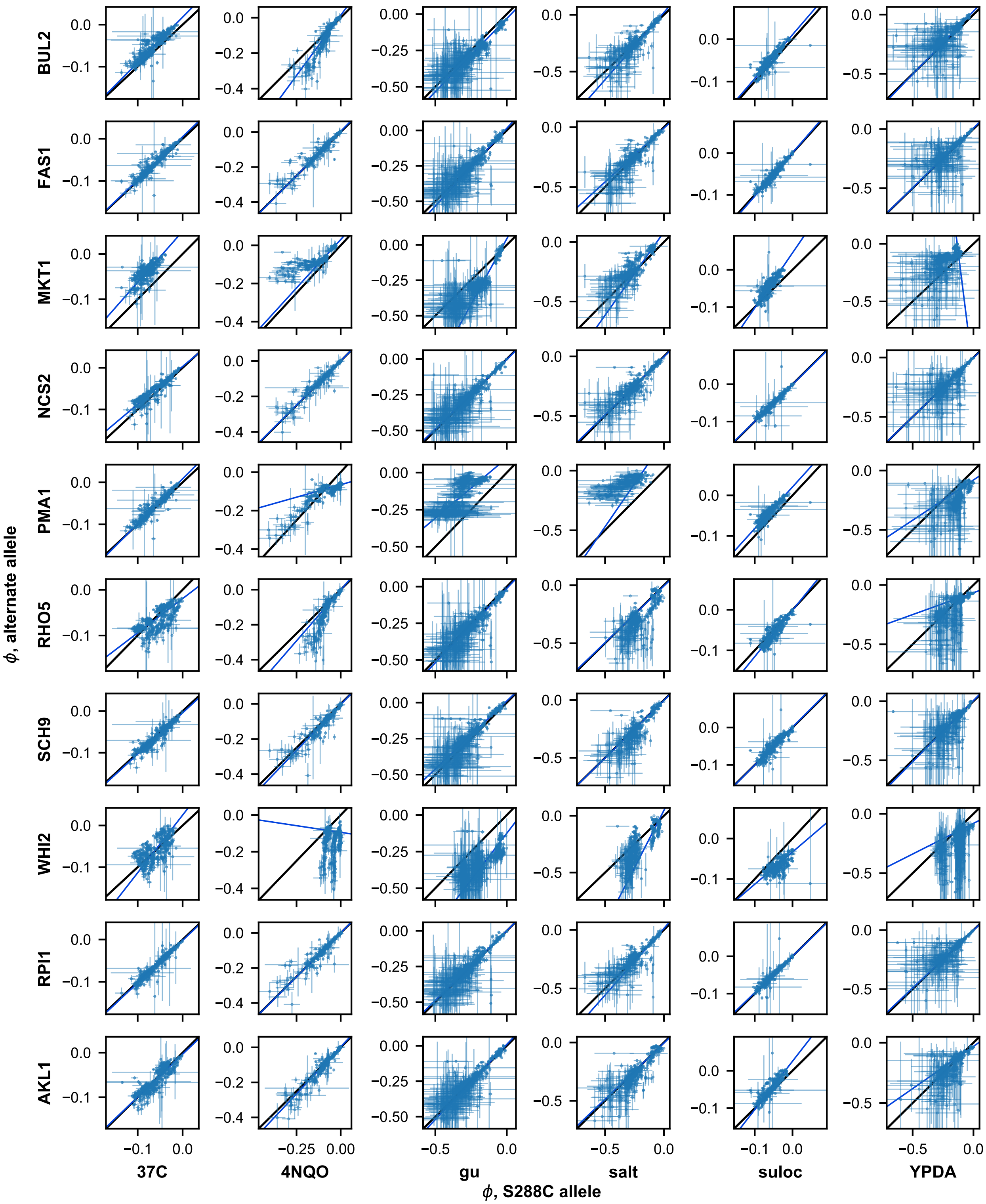

.

Figure S14: Scatterplots of $\phi_{wt}/\phi_{mut}$ for all loci in homozygous form.

had the relevant idiosyncratic interactions, it may be possible to identify the relevant FCT in a smaller landscape. In general, however, because theory argues that it is the accumulation of many random idiosyncratic interactions that produces FCTs, we expect that larger landscapes become more likely to reveal this effect. By random, we mean that idiosyncratic interactions do not obey regular and predictable statistical patterns such as diminishing returns.

In addition to this, another key effect of landscape size is that the total number of genotypes, and hence the total number of fitness measurements, also increases exponentially with the number of mutations in the landscape. This reduces the influence of noise and improves our ability to identify FCTs and the potential effects of idiosyncratic interactions in producing them. This is critical, because linear regression analyses are known to be strongly affected by noise, which can produce outliers: the variance on the slope estimate is (roughly) inversely proportional to the number of data points used in the regression. Since increasing the number of loci considered in fitness landscapes leads to an exponential increase in the total number of data points, we expect that FCTs in significantly smaller landscapes (including landscapes like those examined in previous studies) would therefore be more affected by noise.

To explore these effects of landscape size on the decomposition of fitness-correlated trends (FCTs), we analyzed smaller sub-landscapes from the corresponding subsets of our data. By definition, we cannot disentangle the potential role of idiosyncratic epistasis in creating an FCT in a landscape consisting of only two loci. We therefore constructed landscapes with all possible subsets of three or more of our mutations. For each subset, we analyzed the potential FCT using our decomposition analysis (see Section 6.2). Specifically, for all subsets and all mutations that had evidence of FCT in the full-dataset (i.e., $b \leq 0.9$), we computed the final ratio of sum-squared errors (SSE) between a model with a slope of 1 (this is the idiosyncratic FCT model) and a model with the global initial slope (the global FCT model), after removing all relevant epistatic terms. The idiosyncratic model is supported when this final ratio is below 1. Note that we excluded from this analysis subsets and mutations for which regressions were based on just 1 or 2 points.

We find that, at smaller subset sizes, there is a wide range of final relative fit ratios, indicating that the same mutation can be found to display evidence for either the idiosyncratic model or the global epistasis model driving FCTs. This spread of final SSE ratio can be explained by the random effects of which mutations happen to be represented in each subset, as well as the increased influence of noise on regression and on the inference of coefficients. However, we find that as the subset size increases, the range narrows, with most relative fit ratios dropping below 1 (Figure S15 and Figure S16). This indicates that noise is particularly important in determining whether we can distinguish between the idiosyncratic epistasis model and the global epistasis model, with smaller subsets containing exponentially fewer points and hence far fewer measurements of the fitness effect of mutations (or epistatic terms) with which to perform inference and regression. For our data, with sparse interactions, a landscape of size greater than 8 appears sufficient to provide strong support for the idiosyncratic model (Figure S15 and Figure S16).

To further confirm that noise is the primary driver of evidence towards the global epistasis model (i.e., toward a relative fit ratio $> 1$), we investigated cases where the final relative fit ratio remained above 1 even in our largest fitness landscapes. We found that these have a strong tendency to be mutations in environmental/ploidy combinations with the greatest evidence for noise as determined by the correlation between biological replicates (Figure S17). This suggests that these outstanding cases pointing to global epistasis would be resolved toward the idiosyncratic epistasis explanation with better measurements or with still larger landscapes. We also note that this finding suggests that apparent differences between environments (e.g. with salt and YPDA environments suggesting a larger role for global effects) may simply be an artifact of the inherently noisier fitness measurements in these conditions. These lines of analysis also suggest that previous studies

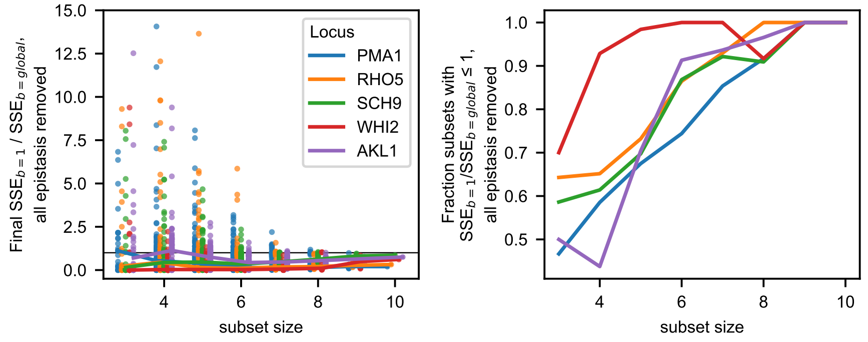

Figure S15: Effects of landscape size on the final SSE ratio (with values less than 1 indicating that FCTs are resolved in terms of idiosyncratic interactions) in 4NQO (haploid). In left panel, each point represents a subset of the full landscape of the corresponding size, with a particular focal mutation (indicated by the legend) having a fitness-correlated slope of $b \leq 0.9$ (polarity adopted such that b is $\leq 1$). The relative fit (sum-squared error, SSE) ratio between regressions with fixed slope of $b=1$ and $b=$global was computed after all epistatic terms were removed. At right, we show the fraction of subsets that have a final (all epistasis removed) relative fit ratio lower than 1 for each mutation, indicating support for the idiosyncratic model of fitness-correlated trends. Not shown are 16 points for which relative fit ratio is greater than 10. Lines show median ratios for each mutation.

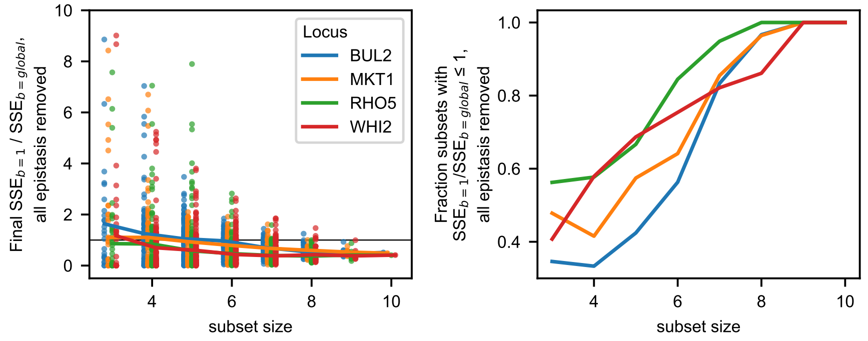

Figure S16: Effects of landscape size on the final SSE ratio (with values less than 1 indicating that FCTs are resolved in terms of idiosyncratic interactions) in 37C (haploid). In left panel, each point represents a subset of the full landscape of the corresponding size, with a particular focal mutation (indicated by the legend) having a fitness-correlated slope of $b \leq 0.9$ (polarity adopted such that b is $\leq 1$). The relative fit (sum-squared error, SSE) ratio between regressions with fixed slope of $b=1$ and $b=$global was computed after all epistatic terms were removed. At right, we show the fraction of subsets that have a final (all epistasis removed) relative fit ratio lower than 1 for each mutation, indicating support for the idiosyncratic model of fitness-correlated trends. Not shown are 15 points for which relative fit ratio is greater than 10. Lines show median ratios for each mutation.

with smaller landscape sizes might not have been able to decompose FCTs as being driven by idiosyncratic epistasis.

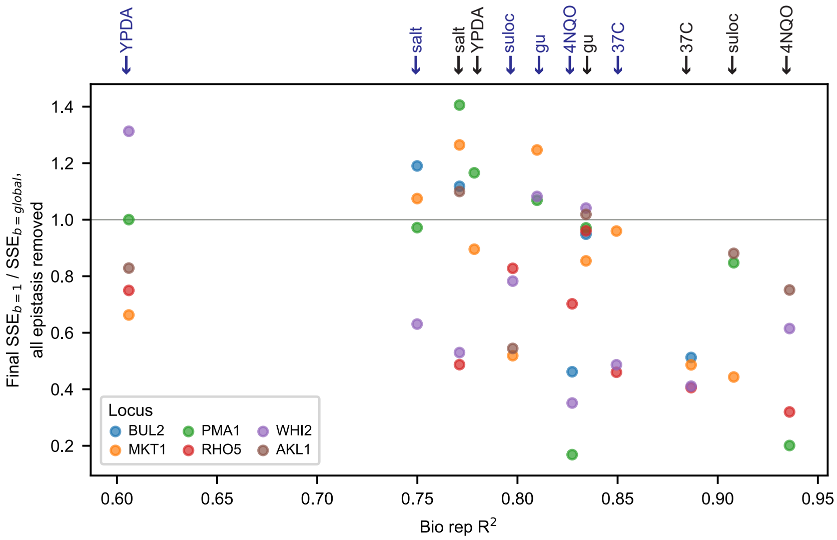

Figure S17: Final relative fit ratio as a function of reproducibility in biological replicates (i.e. the noise in individual fitness measurements). Each point represents the final sum-squared error ratio (i.e. the relative SSE ratio between regressions with a fixed slope of $b$=1 and $b$=global) for a given focal mutation (as indicated in legend) and environment (as indicated by arrows above, with haploids in black and homozygous diploids in blue). Note that SSE ratios greater than 1, which correspond to evidence for global epistasis, occur more frequently when the data is noisier. Only loci exhibiting an FCT in at least 3 of the 12 ploidy/environment combinations are presented.

### 7 Captions for Data Tables

#### 7.1 Data Table S1

Primers used in genotyping, as well as search sequences used in parsing genotypes.

#### 7.2 Data Table S2

Barcode to well to genotype map, and measured competitive fitness of each barcode in each ploidy and each environment.

The fitness values provided are joint inferred fitnesses from two technical replicates (two separate fitness assays were performed simultaneously), and the standard error is of the estimate is obtained from the effect of an overdispersed binomial sampling error on this estimate (see Section 4.3 for more details). The estimated starting frequency of the barcode in the fitness assay in each technical replicate is also provided.

The HSL7-SPT7-FRS1 worksheet indicates whether each well was pure for one or the other allele, or considered impure at a stated threshold.

#### 7.3 Data Table S3

Model parameters for each ploidy in each environment. We provide bootstrap 95% confidence intervals for the parameters as well.

[2] Michael Costanzo, Benjamin VanderSluis, Elizabeth N. Koch, Anastasia Baryshnikova, Carles Pons, Guihong Tan, Wen Wang, Matej Usaj, Julia Hanchard, Susan D. Lee, Vicent Pelechano, Erin B. Styles, Maximilian Billmann, Jolanda van Leeuwen, Nydia van Dyk, Zhen-Yuan Lin, Elena Kuzmin, Justin Nelson, Jeff S. Piotrowski, Tharan Srikumar, Sondra Bahr, Yiqun Chen, Raamesh Deshpande, Christoph F. Kurat, Sheena C. Li, Zhijian Li, Mojca Mattiazzi Usaj, Hiroki Okada, Natasha Pascoe, Bryan-Joseph San Luis, Sara Sharifpoor, Emira Shuteriqi, Scott W. Simpkins, Jamie Snider, Harsha Garadi Suresh, Yizhao Tan, Hongwei Zhu, Noel Malod-Dognin, Vuk Janjic, Natasa Przulj, Olga G. Troyanskaya, Igor Stagljar, Tian Xia, Yoshikazu Ohya, Anne-Claude Gingras, Brian Raught, Michael Boutros, Lars M. Steinmetz, Claire L. Moore, Adam P. Rosebrock, Amy A. Caudy, Chad L. Myers, Brenda Andrews, and Charles Boone. A global genetic interaction network maps a wiring diagram of cellular function. *Science*, 353(6306), 2016.

[18] Mireya Plass. Yeast intron structures database. http://regulatorygenomics.upf.edu/Software/Yeast_Introns/index.html.

[19] Carola Engler, Romy Kandzia, and Sylvestre Marillonnet. A one pot, one step, precision cloning method with high throughput capability. *PloS One*, 3(11):e3647, 2008.

[20] Carola Engler, Ramona Gruetzner, Romy Kandzia, and Sylvestre Marillonnet. Golden gate shuffling: A one-pot DNA shuffling method based on type IIs restriction enzymes. *PLOS ONE*, 4(5):e5553–, 2009.

[21] Michael Baym, Sergey Kryazhimskiy, Tami D. Lieberman, Hattie Chung, Michael M. Desai, and Roy Kishony. Inexpensive multiplexed library preparation for megabase-sized genomes. *PloS One*, 10(5):e0128036, 2015.

[22] Milo S. Johnson, Alena Martsul, Sergey Kryazhimskiy, and Michael M. Desai. Higher-fitness yeast genotypes are less robust to deleterious mutations. *Science (New York, N.Y.)*, 366(6464):490–493, October 2019.

[23] Matthew Barnett. regex: Alternative regular expression module, to replace re.

[24] Alex N. Nguyen Ba, Katherine R. Lawrence, Artur Rego-Costa, Shreyas Gopalakrishnan, Daniel Temko, Franziska Michor, and Michael M. Desai. Barcoded Bulk QTL mapping reveals highly polygenic and epistatic architecture of complex traits in yeast. Technical report,

September 2021. Company: Cold Spring Harbor Laboratory Distributor: Cold Spring Harbor Laboratory Label: Cold Spring Harbor Laboratory Section: New Results Type: article.

[25] Peng Zhao and Bin Yu. On model selection consistency of lasso. *J. Mach. Learn. Res.*, 7:2541–2563, 2006.

[26] Jian Yang, S Hong Lee, Michael E Goddard, and Peter M Visscher. GCTA: a tool for genome-wide complex trait analysis. *American journal of human genetics*, 88(1):76–82, 2011.

[27] David Berger and Erik Postma. Biased estimates of diminishing-returns epistasis? Empirical evidence revisited. *Genetics*, 198(4):1417–1420, December 2014.

[28] Derek York, Norman M. Evensen, Margarita López Martınez, and Jonás De Basabe Delgado. Unified equations for the slope, intercept, and standard errors of the best straight line. *American Journal of Physics*, 72(3):367–375, February 2004. Publisher: American Association of Physics Teachers.
